# A stable, long-term cortical signature underlying consistent behavior

**DOI:** 10.1101/447441

**Authors:** Juan A. Gallego, Matthew G. Perich, Raeed H. Chowdhury, Sara A. Solla, Lee E. Miller

## Abstract

Animals readily execute learned motor behaviors in a consistent manner over long periods of time, yet similarly stable neural correlates remained elusive up to now. How does the cortex achieve this stable control? Using the sensorimotor system as a model of cortical processing, we investigated the hypothesis that the dynamics of neural latent activity, which capture the dominant co-variation patterns within the neural population, are preserved across time. We recorded from populations of neurons in premotor, primary motor, and somatosensory cortices for up to two years as monkeys performed a reaching task. Intriguingly, despite steady turnover in the recorded neurons, the low-dimensional latent dynamics remained stable. Such stability allowed reliable decoding of behavioral features for the entire timespan, while fixed decoders based on the recorded neural activity degraded substantially. We posit that latent cortical dynamics within the manifold are the fundamental and stable building blocks underlying consistent behavioral execution.

## INTRODUCTION

For learned actions to be executed reliably, the cortex must integrate sensory information, establish a motor plan, and generate appropriate motor outputs to muscles. Animals, including humans, readily perform such behaviors with remarkable consistency, even years after acquiring the skill. How does the brain achieve this stability? Is the process of integration and planning as stable as the behavior itself? Here, we explore these fundamental questions from the perspective of populations of cortical neurons. Recent theoretical and experimental work suggests that neural function may be built on the activation of specific population-wide activity patterns – *neural modes* – rather than on the independent modulation of individual neurons^1–5^. These neural modes are the dominant co-variation patterns within the neural population^1^. In experimental scenarios, the activity of the full neural population within the cortex can only be partially sampled, yet the neural modes can be empirically estimated by applying a dimensionality reduction technique^1,6,7^ such as principal component analysis (PCA) to the recorded activity^8^. The set of neural modes defines a *neural manifold*^1,5,9,10^, a surface that captures most of the variance in the recorded neural activity (Fig. 1). We refer to the time-dependent activation of the neural modes as their *latent dynamics*^7,11,12^. In this framework, the activity of each recorded neuron expresses a weighted combination of the latent dynamics from all the modes (Fig. 1b). The neural modes and their latent dynamics have provided increased understanding of the function of many regions throughout the brain^1,5,13–16^, insights that were not apparent at the level of individual neurons.

**Figure 1.**
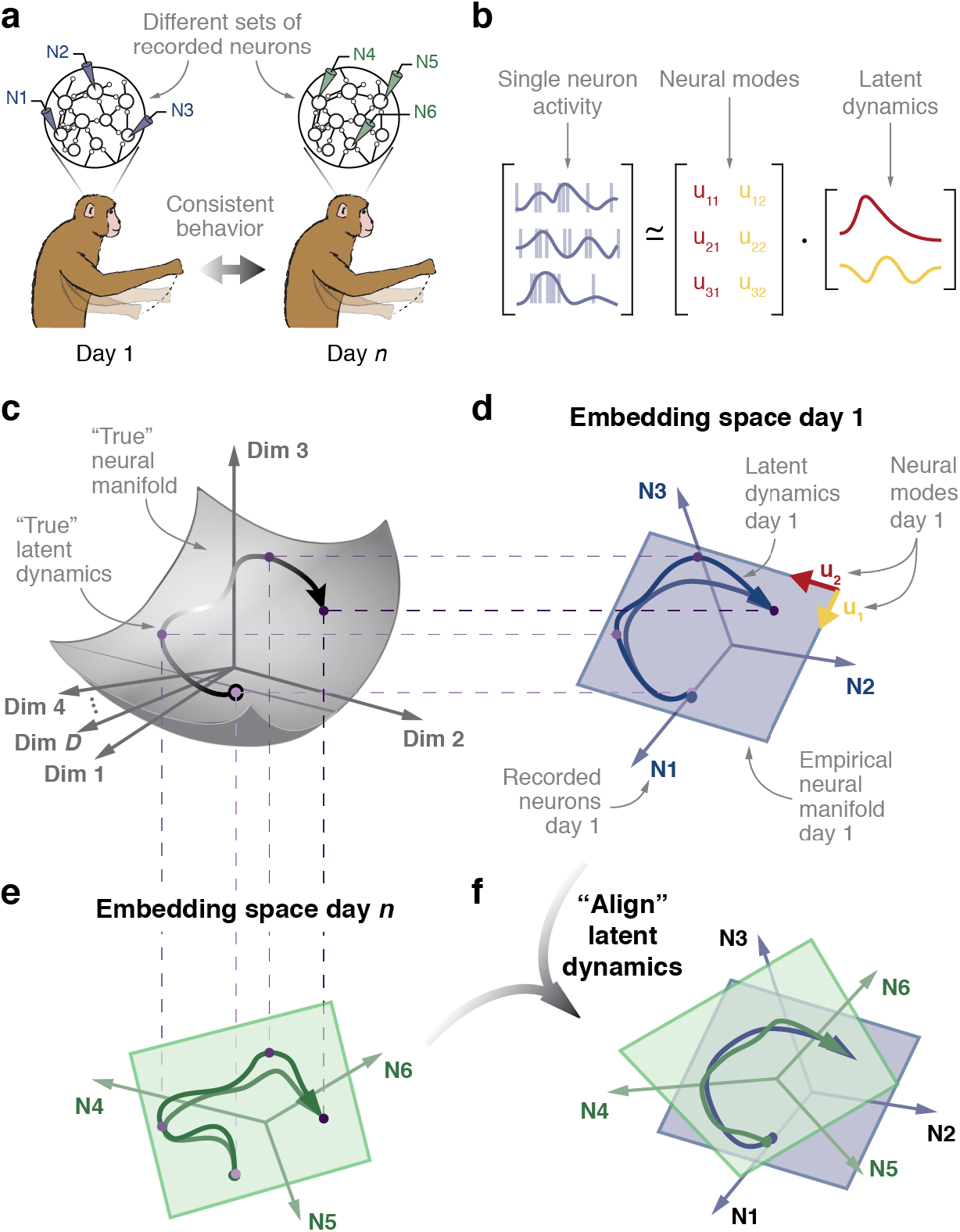
We hypothesize that different movement behaviors are caused by the flexible activation of combinations of neural modes. **(a)** The network connectivity within cortex results in the emergence of neural modes whose combined activation corresponds to specific activity patterns of the individual neurons. **(b)** Neural space for the activity patterns of the three neurons recorded in (a). The time-dependent population activity is represented by the trajectory in black (arrow indicates time direction). This trajectory is mostly confined to a two-dimensional neural manifold (gray plane) spanned by two neural modes (green and blue vectors). **(c)** The activity of each recorded neuron is a weighted combination of the time-varying activation of the neural modes. **(d)** Do neural manifolds for different tasks (show in gray and light purple) have similar orientations? Are the time-varying activations of the neural modes for two tasks (shown in black and purple) similar? These are the two critical questions that test our hypothesis.

We hypothesized that the ability to perform a given behavior in a consistent manner requires that the latent dyna mics underlying the behavior also be stable (Fig. 1). These latent dynamics exist in a relatively low-dimensional manifold that can in principle be estimated using any sufficiently large sample of recorded neurons^7,11^ (Fig. 1c-e). In virtually all studied laboratory tasks, it has sufficed to sample from tens to hundreds of neurons that are modulated by the cortical function being analyzed^7^. However, to quantify the stability of the underlying latent dynamics, the activity of these neurons would have to be recorded over long periods of time, typically months. This need for such stable recordings poses a challenge to current recording techniques such as multielectrode arrays^17,18^. Since each recording session likely samples a somewhat different set of neurons, it is difficult to disentangle differences in the latent dynamics from changes in the recorded neurons and thus evaluate the stability of those dynamics.

Here, we developed a method to examine the stability of the underlying latent dynamics despite these unavoidable changes in the set of neurons recorded using chronically implanted microelectrode arrays. With this method, we addressed the question of stability of latent dynamics, using the sensorimotor system as a model of cortical processing. We recorded the activity of neural populations, approximately one hundred neurons at a time, in each of three different cortical areas: dorsal premotor cortex (PMd), primary motor cortex (M1), and primary somatosensory cortex (S1), as monkeys performed the same reaching behavior. PMd plays a critical role in movement planning, exhibiting strong pre-movement preparatory activity^19^ that can be used to decode the intended movement well before it occurs^20^. M1 is the primary cortical area from which descending output to the spinal cord arrises^21,22^; its activity is tightly coupled to the dynamics of motor execution^23–25^, even during motor adaptation^26^. Lastly, area 2 of S1 receives and integrates both muscle and somatosensory feedback^27,28^, and likely plays a critical role in correcting ongoing movements^29^.

In all three cortical regions, PMd, M1, and S1, we found remarkably stable latent dynamics for up to two years, despite large, on-going turnover of the recorded neurons. The stable latent dynamics, once identified, allowed for the prediction of various behavioral features, using models whose parameters were fixed throughout the entirety of these long timespans. Interestingly, these models predicted behavioral features virtually as accurately as similar models that were trained and tested within the same day. Therefore, we have identified a neural correlate of stable behavior: the low-dimensional latent dynamics of cortex, which underlie the full-dimensional neural population activity. Given that our results hold for three cortical regions involved in different sensorimotor functions, we posit that analogous stable latent dynamics may be used broadly throughout cortex when performing a variety of learned functions, from stimulus recognition to complex cognitive processes.

## RESULTS

### Hypothesis and approach

We studied the stability of the latent dynamics within the neural manifold, and their relationship with a behavior that was performed consistently over many days (Fig 1a). To identify the neural modes, we represented the activity of each recorded neuron along one axis in a high-dimensional embedding neural space^1,8,9^. In the toy example in Fig. 1d, the number of recorded neurons, and thus the dimensionality of the neural space, is three. The neural modes can be computed using any dimensionality reduction method that identifies patterns of neural covariation^8^; here we used principal component analysis (PCA) applied to smoothed firing rates^8^ (see Methods). Mathematically, the PC axes are the neural modes that span the neural manifold^1,5,9,10^ (plane in Fig. 1d).

By projecting the recorded neural activity into each neural mode, we can calculate an empirical estimate of the “true” latent dynamics throughout the cortex^7^ (Fig. 1b,d). Changes in the neurons recorded over days necessarily cause a change in the axes that empirically define the embedding neural space (compare Fig. 1d to 1e), along with a corresponding change in the empirically estimated manifold and latent dynamics. These changes do not necessarily imply a change in the true latent dynamics governing cortical function; they may instead simply represent a different projection of the true stable dynamics onto a new embedding neural space (Fig. 1c-e). Since the projection from the full neural space to the empirical neural space of recorded neurons is linear, and PCA identifies a linear empirical neural manifold within the empirical neural space. Thus, if the true latent dynamics during repeated task execution were indeed stable, a simple linear transformation should be sufficient to align the projection of the latent dynamics within different empirical neural spaces (Fig. 1f). This linear transformation may be a combination of global translation, rotation, scaling, shearing, and reflection. If the true latent dynamics were to change fundamentally across days, e.g., due to a change in the intrinsic dynamics of the network^30^, no linear transformation would be able to align the dynamics across different empirical manifolds. We have developed methods based on canonical correlation analysis (see Methods) to compensate for changes in recorded neurons and to compare the empirical latent dynamics over time; we will refer to this process as one of “aligning” the latent dynamics. In the following sections, we test the hypothesis that consistent behavior is associated with stable latent dynamics by analyzing neural population activity recorded in sensorimotor cortex over weeks to years.

### Behavior

We trained six monkeys (Monkeys C, M, T, J, H, P) to perform a center-out reaching task with an instructed delay period, using a planar manipulandum (Methods; Fig. 2a). The monkey started each trial by holding in the central target, after which one of eight outer targets was presented (Fig. 2b). Following a variable delay period during which the target continued to be visible, a go cue was given, and the monkey had to move the manipulandum to the presented target to receive a liquid reward. Monkey C initially performed the task using the left hand; later, he used the right hand during experiments with multielectrode arrays implanted in the opposite hemisphere (C_L_ and C_R_). Across monkeys, the time between recordings spanned from ~20 to ~750 days (see Table 1), allowing us to study the stability of latent dynamics over long periods of time.

**Figure 2.**
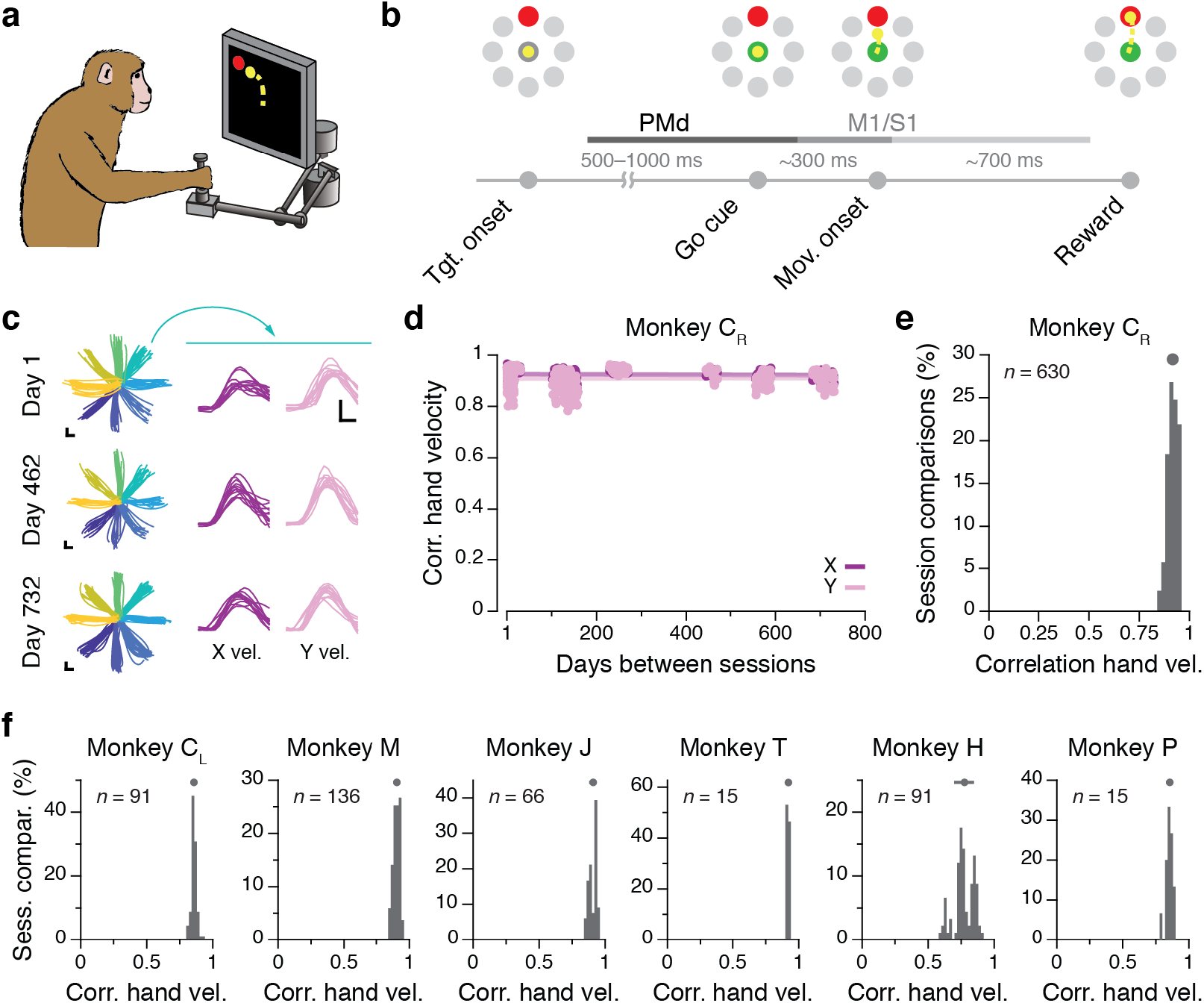
Task and repeatability of behavior. **(a)** Monkeys performed an instructed-delay reaching task using a planar manipulandum. **(b)** Schematic of the task, indicating the approximate windows used for analysis; these varied across cortical areas (PMd, M1, and S1). **(c)** Left: Example hand trajectories for three days spanning 731 days from Monkey C_R_. Each trace is an individual trial; traces are color-coded based on target location. Right: Example X and Y hand velocity traces for all reaches to the upper-right target on each of the three days. **(d)** Correlation between direction-matched single trial X and Y hand velocities across all pairs of days from Monkey C_R_ (single dots: individual pair of days; lines: linear fits). **(e)** Distribution of across-day hand velocity correlations for all pairs of days from Monkey C_R_. Top error bars: mean ± s.d. **(f)** Same as (e), for data from the other six implants.

**Table 1.** Database of neural and behavioral recordings

| Monkey | Age during experiment (years) | Implant site | Number of recording sessions | Range of days | Minimum number of rewarded reaches per target and session |
| --- | --- | --- | --- | --- | --- |
| M | 6-7 | Right PMd | 17 | 508 | 17 |
|  |  | Right M1 | 18 | 508 | 17 |
| T | 10 | Right PMd | 6 | 21 | 17 |
| C<sub>R</sub> | 6-8 | Right M1 | 36 | 731 | 12 |
| C<sub>L</sub> |  | Left PMd | 14 | 42 | 10 |
|  |  | Left M1 | 14 | 42 | 10 |
| J | 19 | Left M1 | 12 | 71 | 16 |
| P | 8 | Left S1 | 6 | 28 | 10 |
| H | 8 | Left S1 | 14 | 44 | 12 |

The behavioral performance of all monkeys was consistent across days, as exemplified by the hand trajectories in Fig. 2c. To quantify stability, we computed the correlation between the X and Y hand velocities across single trials for a given target, for all pairs of days (Fig. 2d). In all of the cases, these correlations were large (mean >0.77, Fig 2d-f; Fig. S1 replicates Fig. 2d for the other monkeys).

### Changes in single neuron recordings across days

We studied the neural basis for the consistent behavior shown in Fig. 2 using 96-channel microelectrode arrays chronically implanted in the arm areas of three different regions of cortex (Methods); the approximate location of the implanted arrays are shown in Fig. 3a. Monkeys C_L_ and M had dual implants in PMd and M1; Monkeys J and C_R_ had an implant in M1; Monkey T had an implant in PMd; and Monkeys H and P had an implant in Area 2 of S1. We manually spike-sorted the neural recordings from Monkeys C, M, and T to compare whether the same neurons were recorded across days (Methods; Fig. S2 shows example neurons). As illustrated by the M1 dataset from of Monkey C_L_ (Fig. 3b), single neuron activity changed quite dramatically over 15 days. In addition to some variation in the behavior itself^18^, it is likely that many of these changes can be explained by turnover in the neurons recorded on each electrode (Figure 3c). To quantify the turnover effect, we tracked the stability of both firing rate statistics and waveform shape of each neuron^31,32^ (Fig 3c, bottom, shows one example). Across all pairs of days for this dataset, the percentage of matched neurons decreased to < 30 % after 25 days (Fig. 2d; Fig S3 shows similar results for six other sets of recordings from M1 and PMd).

**Figure 3.**
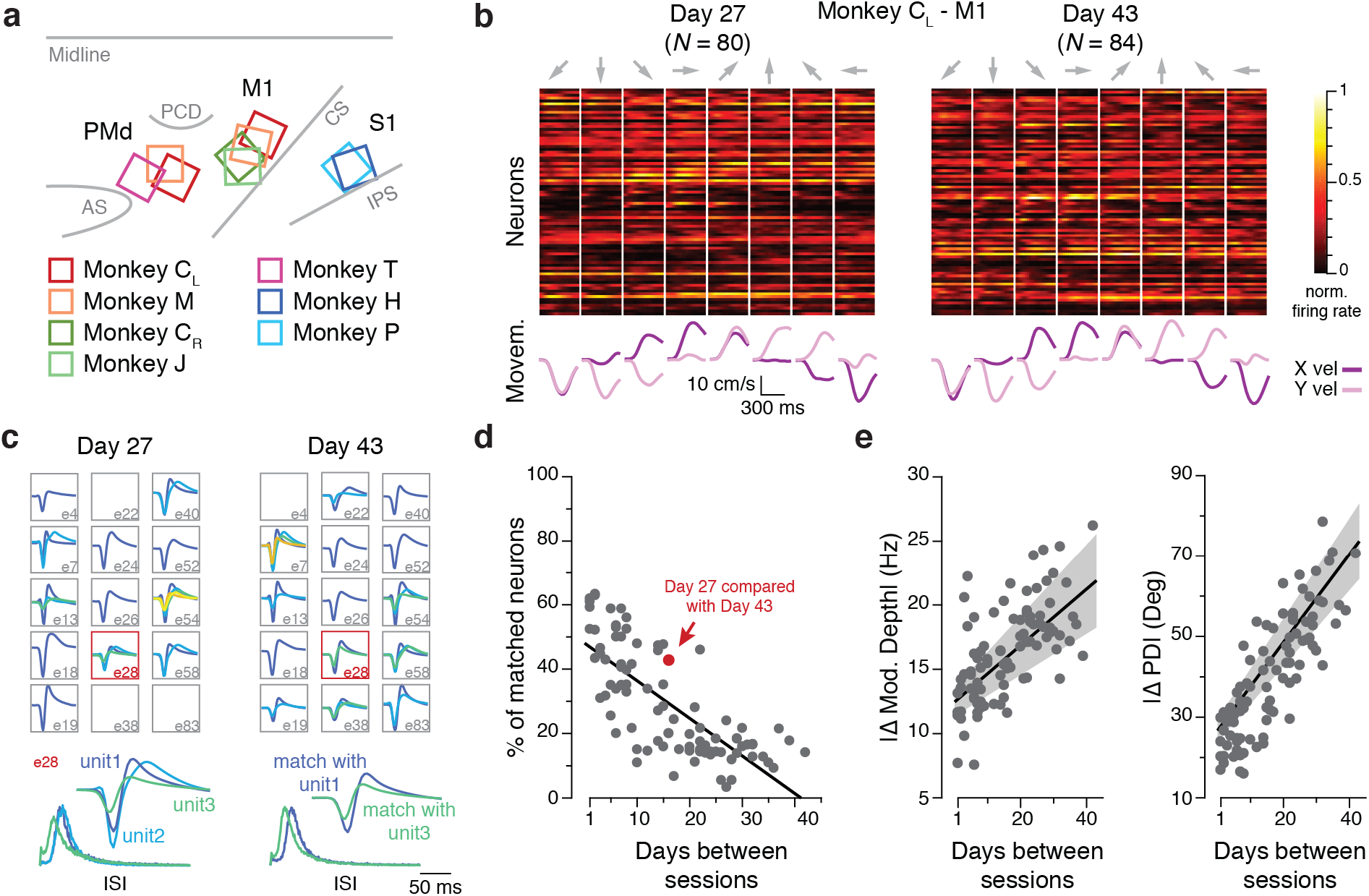
Neural recording and functional tuning stability. **(a)** Approximate location of all nine arrays for the seven implants; each monkey is represented by a different color (legend). AS, arcuate sulcus; PCD, precentral dimple; CS, central sulcus; IPS, intraparietal sulcus. **(b)** Peristimulus time histograms for all sorted neurons identified on Day 27 and Day 43 from Monkey C_L_ (top; each neuron shown in a different row) and corresponding hand velocity (bottom). Each column represents the average of all trials to each of the eight reach directions (indicated by the arrows above each column). Note the substantial changes in the activity of the recorded neurons, reflected in altered firing rates and spatial tuning, despite the consistency of the behavior. **(c)** Example sorted neurons for these two datasets; note the large apparent turnover after 15 days. Bottom inset: example action potential waveforms and inter-spike interval (ISI) histograms for two neurons that were matched across days. **(d)** Percentage of individual neurons matched across all pairs of days from Monkey C_L_; matching based on the same methods as in (c). **(e)** Change in modulation depth (left), and preferred direction (right) of standard cosine tuning fits to multiunit activity. Data for all pairs of days from Monkey C_L_. Line and shaded areas: mean ± s.e.m.

In addition to tracking characteristics of well discriminated single neurons, we assessed neural turnover by fitting tuning curves relating neural activity to the reach direction (Methods). For this and all subsequent analyses, we used the multiunit activity (threshold crossings) recorded on each electrode rather than sorted neurons, to incorporate as many neural events as possible while preserving our ability to reconstruct the latent dynamics^33^. We found that both modulation depth and preferred direction changed progressively over time, as shown in Fig. 3e for one representative dataset. There were similar progressive changes for all monkeys and cortical regions (Fig. S4). Combined, these analyses indicate that the set of recorded neurons changed substantially across days, with consequent changes in estimates of spatial tuning from the array recordings. Our hypothesis predicts that, despite these changes in recorded neurons, the underlying latent dynamics of the full neural population will be stable.

### Primary motor cortex during movement control

We now investigate the hypothesis that stable latent dynamics within the cortex underlie the generation of consistent motor behavior, starting with the analysis of M1 activity during movement execution. We applied our method to “align” the latent dynamics across the manifolds for different days, even as the number and identity of recorded neurons changed. We used single trial neural data to compute the neural manifold and the latent dynamics within it using PCA separately for each day (Methods). The dimensionality of the manifold for each brain region (M1: 10; PMd: 15; S1: 8) was selected based on previous studies^12,15^. Using canonical correlation analysis^12,30,34^ (CCA), we found the linear transformations that make the latent dynamics from day *n* maximally correlated to those of day one (Methods). These transformations compensated for the changes in the recorded population of neurons caused by turnover. As described above, if the true latent dynamics during repeated behavior were indeed stable, the trajectories of the empirically estimated latent dynamics would be very similar after alignment (Fig. 1f): the leading canonical correlations (CCs) would approach a value of one. On the contrary, if repeated behavior did not result from stable underlying latent dynamics within the brain, the trajectories would be very different even after attempted alignment, and all the resulting CCs would be low.

The trajectories described by the empirical latent dynamics for datasets separated by 31 days were indeed very different (Fig. 4a), presumably reflecting the observed changes in recorded neurons (Fig. 3, S2, S3). However, these trajectories became quite similar after alignment with CCA (Fig. 4b).

**Figure 4.**
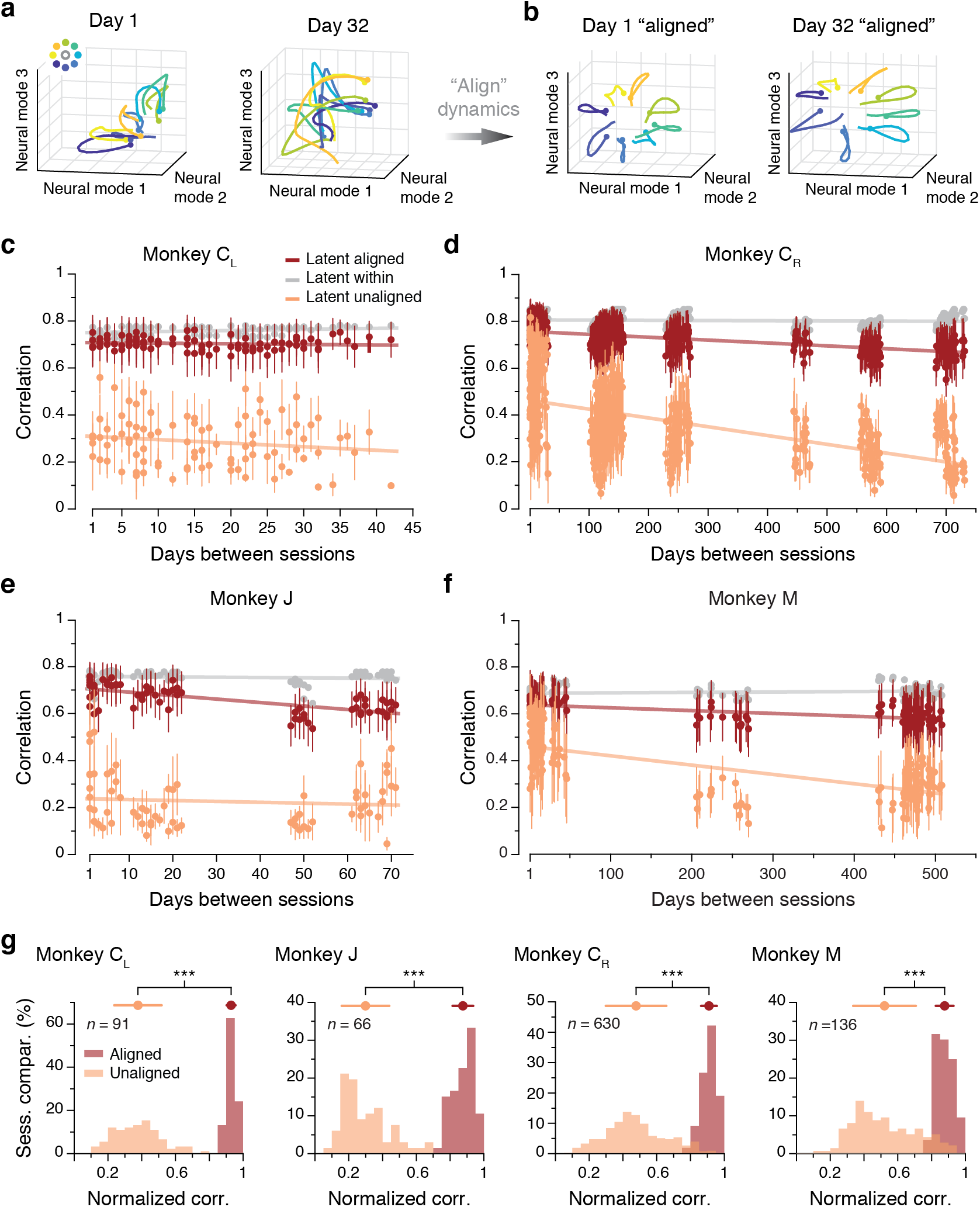
Stability of M1 latent dynamics over time. Example **(a)** “unaligned” and **(b)** “aligned” latent trajectories for two example days from Monkey C_L_. Note the similarity between the latent trajectories to acquire each target after alignment. Data averaged over all trials for a fixed target only for visualization purposes. **(c)** The correlation of the M1 latent dynamics averaged over the top four neural modes across all pairs of days from Monkey C_L_ (single dots: pairs of days; lines: linear fits). The aligned latent dynamics (red) maintained a higher correlation across days than the unaligned dynamics (orange), and were almost as correlated across different days as the latent dynamics across different blocks of trials from the same day (gray). **(d)-(f)** Same as (c) for the other three M1 implants. **(g)** Normalized similarity of the aligned and unaligned M1 latent dynamics during movement execution for each monkey (each shown in a different panel). The mean normalized similarity of the aligned latent dynamics across different days is close to 1; this indicates that their canonical correlations are almost as strong as the canonical correlations of the latent dynamics across two blocks of trials from the same day. *** indicates ***P*** < 0.001 per Wilcoxon rank-sum test. Error bars: mean ± s.d.

This observation held across all pairwise combinations of days for this monkey; the aligned empirical latent dynamics remained stable for over 40 days, the full length of time we were able to monitor for this monkey (Fig. 4c).

To better interpret the magnitude of this across-day stability, we compared it to the stability across random blocks of trials within each day, meant to provide an upper bound on the achievable CCs (Methods). For this monkey, the across-day CCs were virtually identical to the within-day CCs (red and gray traces in Fig. 4c). To summarize these results, we computed a *normalized similarity*: the ratio of the across-day CCs to the corresponding within-day CCs. For this dataset, the normalized similarity of the latent dynamics after alignment with CCA was 0.93 ± 0.03 (mean ± s.d.; Fig. 4g). The normalized similarity without alignment was much lower (0.38 ± 0.14; Fig. 4g). We obtained similar results for all M1 datasets, comprising four implants from three different monkeys (Fig. 4d-f,g); this indicates that M1 latent dynamics during repeated movement generation are stable for very long periods of time.

Many neuroscientific studies have attempted to understand the information encoded within neural populations by predicting relevant behavioral features using neural activity. This is of particular interest within the field of brain-computer interfaces that seek to map motor cortical activity onto control signals for prostheses or paralyzed limbs^23,25,35–37^ - the limited stability of these predictive models over time has been an ongoing source of concern^18,38^. Consequently, we asked how accurately a linear model trained to predict hand kinematics based on latent dynamics from one day would perform on aligned latent dynamics from a different day (Fig. 5a; Methods). Fig. 5b shows hand velocity reliably predicted 16 days later. This performance was almost as good as that of a model trained and tested on the same day, and much better than that of a model based directly on the recorded multiunit neural activity (Fig. 5b) or on the unaligned latent dynamics (Fig. S5a,b). This observation held for all pairwise comparisons of days for this dataset (Fig. 5c). To summarize these comparisons, we computed a *normalized predictive accuracy*: the ratio of the across-day R^2^ to the within-day R^2^. For this representative example, the normalized predictive accuracy across all pairs of days averaged 0.96, indicating that the aligned latent dynamics provide nearly the same behavioral predictive ability as the neural activity recorded on the same day the model was built (Fig. 5g, left). However, the normalized predictive accuracy of models based directly on the recorded neural activity degraded steadily with time after the model was built, to R^2^ values as low as 0.13 (Fig. 5c; mean R^2^ of 0.60). We replicated these results on datasets from Monkeys C_R_, J, and M (Fig. 5d-f,g); for monkeys C_R_ and M we could even predict hand velocity accurately two years and one and a half years later, respectively (Fig. 5d,f). Thus, the relationship between recorded neural activity and behavior changes day to day, and the predictive ability of a model trained once and held fixed quickly deteriorates over time. However, the low-dimensional latent dynamics can be aligned to maintain the good predictive ability of a fixed model. Models based on the aligned latent dynamics predict behavioral features almost as well as models trained on same day neural recordings.

**Figure 5.**
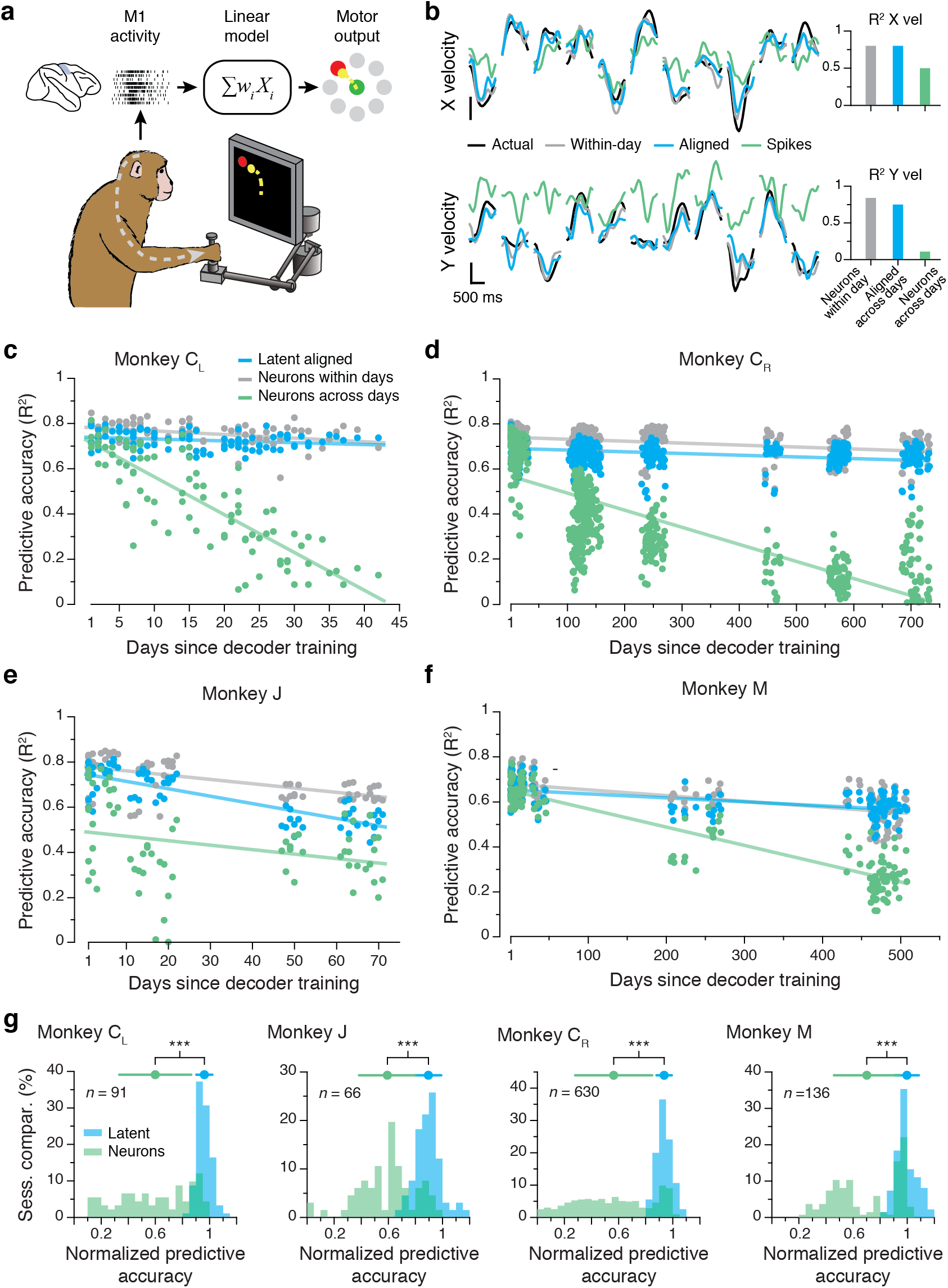
Stable decoding of movement kinematics based on the aligned latent dynamics. **(a)** We trained linear models to predict movement kinematics based on different types of inputs. **(b)** Example X and Y velocity predictions for two recordings made 16 days apart. Predictions based on the aligned latent dynamics were almost as good as predictions based on the recorded neural activity when trained and tested on the same day (bars on the left show the R^2^ for the entire day). **(c)** Predictive accuracy for models trained and tested on all pairwise combinations of days (single dots: pairs of days; lines: linear fits). Models based on the aligned latent dynamics (green) performed almost as well as models trained and tested on the same day (gray), and much better than models trained on the recorded neural activity when tested across days (blue). **(d)-(f)** Same as (c) for the other three M1 implants. **(g)** Normalized accuracy for models based on the aligned latent dynamics (green) and the recorded neural activity (blue) when tested on a different day. Each panel shows one monkey; each data point is one pairwise comparison between days. The mean normalized predictive accuracies of the aligned latent models are close to 1; this indicates that they as predictive about behavior as models trained on the recorded neural activity for each day and tested on the same day. *** indicates ***P*** < 0.001 per Wilcoxon rank-sum test. Error bars: mean ± s.d.

We performed two controls to verify the robustness of our results. Although our alignment analysis is based on a choice of manifold dimensionality, our stability results were unchanged when the analyses were repeated for a wide range of manifold dimensionalities (Fig. S5c-e). Additionally, previous work has shown that empirically estimated latent dynamics are quite similar whether the manifold is obtained using sorted neurons or multiunits^33^. Here we generalize this result to longterm stability, showing for three monkeys in Fig. S5f,g that our main result also holds when using sorted neurons to identify the manifold.

### Dorsal premotor cortex during movement planning

We have shown M1 latent dynamics during repeated, consistent behavior to be stable for very long periods of time, allowing for stable decoding of movement kinematics. As M1 is primarily involved in the execution aspect of movement generation, we asked whether a similar stability principle might apply to PMd activity during motor planning (Fig. 6a). Pre-movement planning activity in PMd captures many features of the subsequent behavior, including reaction time^39^, speed^40^, and even the ability to learn^15^. We thus expected that consistent movements would be preceded by motor planning dynamics that are stable over time. We tested the stability of PMd latent dynamics during planning using the same CCA alignment procedure we used for M1. As was the case for M1, despite changes in recorded neurons (example in Fig. S6a; see also Fig. S3,4), PMd latent dynamics were stable over weeks and even months; Fig. 6c and Fig. S6 show data for three different monkeys. The stability of the aligned latent dynamics across days was nearly indistinguishable from within-day stability (Fig. 6c, red and gray, and Fig. 6d, red). Without alignment, the leading latent dynamics were quite different across days (Fig. 6c, orange, and Fig. 6d).

**Figure 6.**
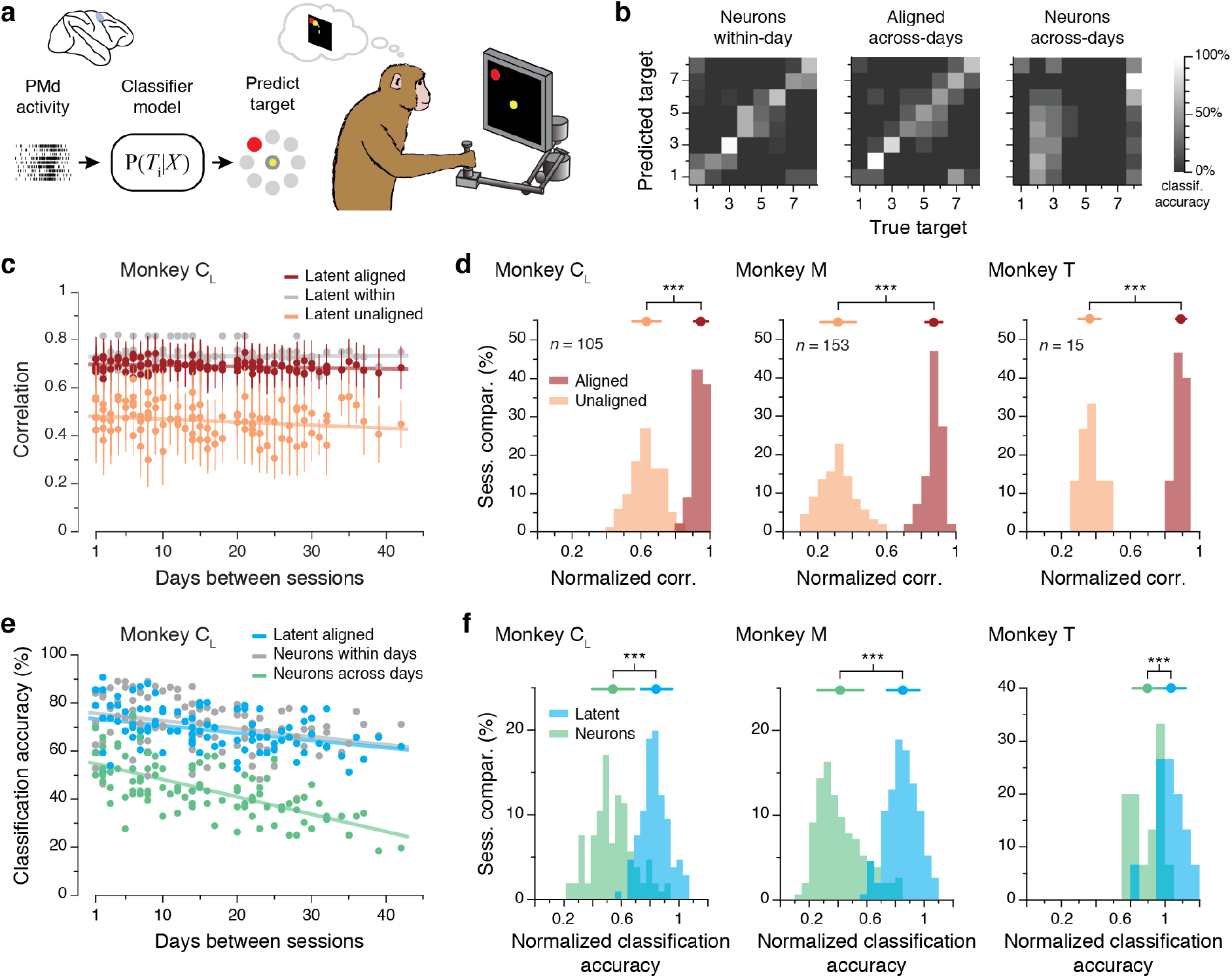
Stability of PMd latent dynamics during movement planning. **(a)** We trained classifier models to predict the intended target based on neural activity, either recorded or latent. **(b)** Example confusion matrix showing classification performance. Within-day models (left) performed quite accurately (73% correct). The performance of across-day models (middle) based on the aligned latent dynamics was well above chance (54%), while models based on recorded neural activity (right) performed quite poorly (23%) when trained and tested on different days. Color bar: classification accuracy. Data from Days 27 and 43 from Monkey C_L_. **(c)** Correlation of the PMd latent dynamics averaged over the top four neural modes across all pairs of days from Monkey C_R_ (single dots: pairs of days; lines: linear fits). The aligned latent dynamics (red) are more highly correlated across days than unaligned dynamics (orange), and almost as highly correlated as the latent dynamics from different blocks of trials from the same day (gray). **(d)** Normalized similarity of the aligned and unaligned PMd latent dynamics during movement planning for each monkey (each shown in a different panel). The mean normalized similarity of the latent dynamics from two different days is close to 1, indicating that they are almost as correlated as the latent dynamics across two blocks of trials from the same day. *** indicates ***P*** < 0.001 per Wilcoxon rank-sum test. **(e)** Classification accuracy for models trained and tested on all different pairs of days. Models based on the aligned latent dynamics (green) performed almost as well as models trained and tested on the same day (gray), and much better than models trained on recorded neural activity and tested on different days (blue). **(f)** Normalized predictive accuracy for models based on the aligned latent dynamics (green) and recorded neural activity (blue) tested on a different day. Each panel shows one monkey; each data point is one pairwise comparison between days. The mean normalized accuracies of the aligned models are close to and even above 1, indicating that they provide almost as much predictive ability about behavior as models trained on neural activity recorded on the same day. *** indicates ***P*** < 0.001 per Wilcoxon rank-sum test. Top error bars: mean ± s.d.

We next tested whether these stable latent dynamics in PMd could be used reliably to predict behavior. Previous work has shown that it is possible to determine the intended movement from the PMd planning activity before it occurs^41^. We used naïve Bayesian classifiers^20^, trained on either the full recorded neural activity or the latent dynamics during the instructed delay period, to predict the upcoming reach target (Fig. 6a; Methods). We asked whether these models would perform well when used days after training, comparing the across-day performance to that of models trained within the test day. Sixteen days after training, the accuracy of models based on aligned latent dynamics remained close to that of models based on neural activity recorded within the same day (Fig. 6b). In general, the accuracy of the across-day models based on aligned latent dynamics was virtually identical to that of within-day models based on recorded neural activity (Fig 6e). The *normalized classification accuracy*, defined as the ratio of the across-day accuracy to the within-day accuracy, shows that alignment of latent dynamics allowed the across-day models to predict nearly as well as the within-day models (Fig. 6f). These results were replicated using PMd datasets from two other monkeys (Fig. 6f, Fig. S6), showing that even models trained up to ~500 days earlier yielded results comparable to a model trained on the same day (Fig. S6c). Comparison of this performance to the steady, progressive performance decline of fixed models based on recorded neural activity (Fig. 6e, Fig S6c) provides further evidence in support of our hypothesis that stable latent dynamics underlie consistent behavior.

### Primary sensory cortex during feedback control

An important aspect of motor control is the integration of sensory feedback. While we receive feedback about our movements through many different modalities, one of the most critical sensations for motor control is proprioception, our sense of body positioning and movement. Studies of patients with impaired proprioception show that this particular kind of feedback is crucial for generating coordinated movements^42–44^. Here, we examine area 2 of primary somatosensory cortex (S1), a proprioceptive part of the brain that integrates feedback from cutaneous and muscle receptors and also shares connections with the motor cortex^29,45–49^. Given the stability of latent dynamics during planning and execution in PMd and M1, we anticipated seeing similar stability in the somatosensory activity of S1. We tested this hypothesis by analyzing data from S1 during reaching (Fig. 7a). After alignment, latent dynamics in S1 were stable for the full ~30 and ~45 day periods we tested, respectively, for both Monkey P and Monkey H (Fig 7b, Fig S7b). As with the other cortical regions, the normalized stability of the aligned latent dynamics was near the upper bounds set by the within-day correlations of latent dynamics for both monkeys (Fig. 7d).

**Figure 7.**
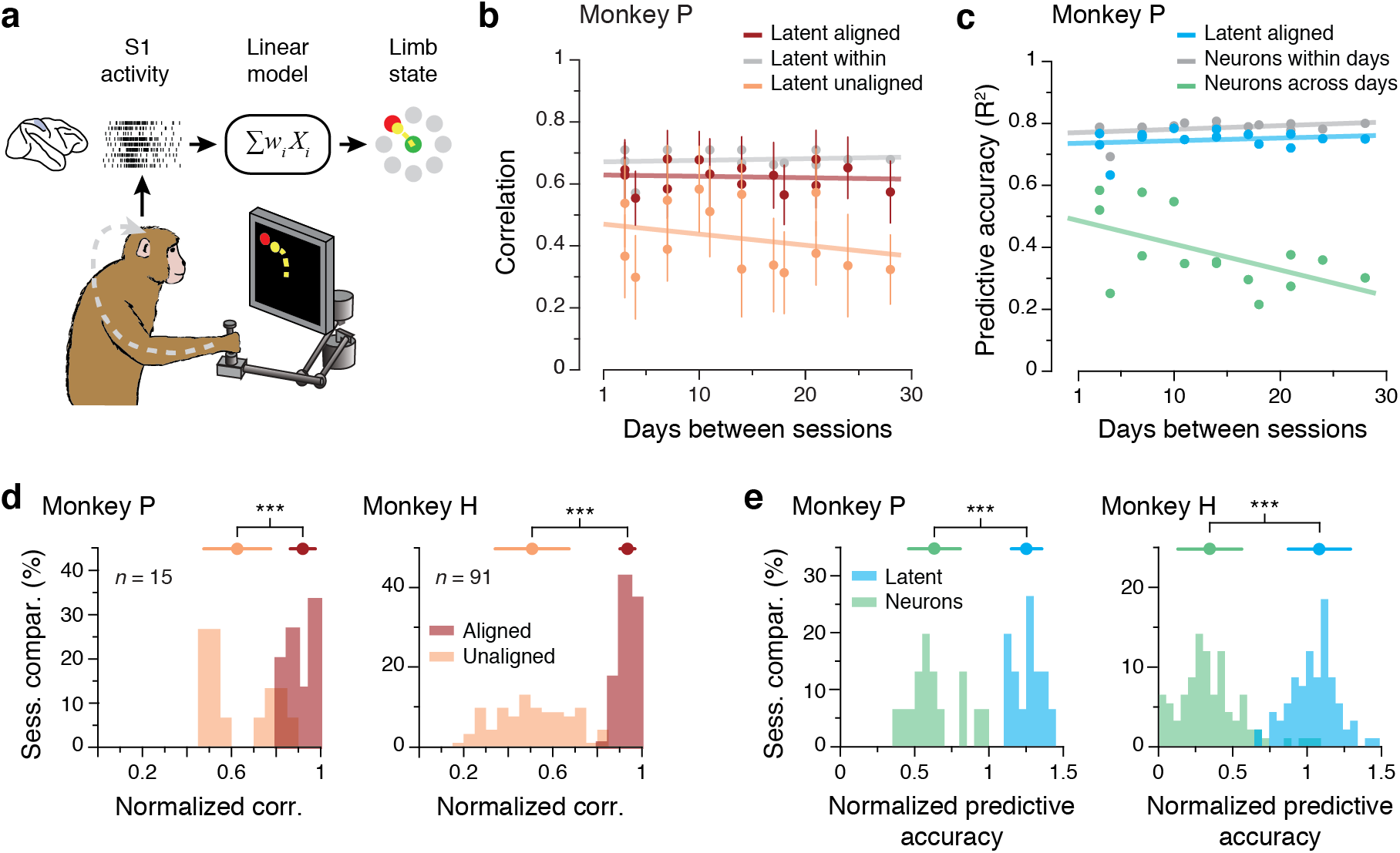
Stability of S1 latent dynamics during feedback control. **(a)** Models trained to predict movement kinematics based on different types of inputs. **(b)** Correlation of the S1 latent dynamics averaged over the top four neural modes across all pairs of days from Monkey P (single dots: pairs of days; lines: linear fits). The aligned latent dynamics (red) are much more correlated across days than the unaligned dynamics (orange), and almost as correlated as the latent dynamics from different blocks of trials from the same day (gray). **(c)** Predictive accuracy for models trained and tested on all different pairs of days. Models based on the aligned latent dynamics (green) performed almost as well as models trained and tested on the same day (gray), and much better than models trained on recorded neural activity when tested across days (blue). **(d)** Normalized similarity of the aligned (red) and unaligned (orange) S1 latent dynamics during movement execution. Each panel shows one monkey; each data point is one pairwise comparison between days. The mean normalized similarity of the latent dynamics from two days is close to 1. This indicates that they are almost as similar as the latent dynamics across two blocks of trials from the same day. *** indicates ***P*** < 0.001 per Wilcoxon rank-sum test. **(e)** Normalized predictive accuracy for models based on the aligned latent dynamics (green) and recorded neural activity (blue) when tested on a different day. Each panel shows one monkey: each data point is one pairwise comparison between days. The mean normalized accuracies of the aligned models are close to 1, indicating that they provide almost as much predictive ability about the behavior as models trained on neural activity recorded on the same day. *** indicates ***P*** < 0.001 per Wilcoxon rank-sum test. Top error bars: mean ± s.d.

S1 encodes rich information about the state of the limb and its evolution over time^29^, which allows for quite accurate decoding of movement kinematics^29^. We tested whether the relationship between aligned latent dynamics and hand velocity was stable over time using models similar to those used for M1 (Fig 7a; Methods). Models based on the aligned latent dynamics predicted movement reliably over time for both monkeys (Fig. 7c, Fig. S7c). As was the case for the motor cortical regions, the accuracy of these predictions was similar to that of models trained and tested on the same day (Fig. 7e). In contrast, the performance of models based on recorded neural activity degraded quickly after a few days (Fig. 7c, Fig. S7c). Taken together, these results suggest that stable behavior is associated with stable latent dynamics throughout sensorimotor cortex, as predicted by our hypothesis.

## DISCUSSION

Once learned, behaviors can be readily executed with accuracy and consistency. How the brain achieves this behavioral stability is still an open question. Existing readouts of neural activity have been fraught with the problem of ever-changing neurons making it extremely difficult to answer questions about the long-term stability of cortical dynamics. Here we have shown that repeated execution of a given behavior is accompanied by stable latent dynamics in several sensorimotor areas of cortex. These stable latent dynamics are associated with all three main aspects of movement generation^50^: planning the upcoming movement, controlling its execution, and integrating somatosensory information. Moreover, the stabilized latent dynamics maintained a fixed relationship with the behavior; remarkably, models based on the stabilized latent dynamics predict behavior up to two years after training almost as well as models built within the same day. Our results show surprising stability in the latent dynamics underlying the consistent execution of the same behavior over long periods of time.

One crucial interpretational point is that we have studied the neural dynamics underlying the planning and execution of one stereotyped reaching behavior. The corresponding neural dynamics are intimately and necessarily linked to the behavior that results. A different behavior, such as shaking a cocktail shaker, would require its own dynamics. A comparative analysis of the neural dynamics corresponding to these two different behaviors performed in succession might reveal some similarities; however, they most likely would differ by much more than observed here when monitoring the same behavior over time. The extent to which latent dynamics are shared between behaviors is an area of active investigation. A recent study from our lab has shown both differences and similarities in M1 latent dynamics when comparing different but related wrist movement tasks^12^. Another study in mice has shown even greater differences in M1 dynamics between forepaw reaching and quadrupedal locomotion^51^. The present results show that our empirical estimates of the latent dynamics can vary greatly, yet when a specific behavior is consistently repeated, the true latent dynamics are stable over months to years. The differences across days arise from projecting the true latent space onto empirical neural spaces that keep changing; these differences can be compensated for through the simple linear transformations employed by CCA.

Given the ubiquity of linear population-level analyses such as PCA in modern neuroscience experiments, our fundamental observation has important implications for understanding how neural populations throughout the brain consistently perform behaviorally relevant functions.

## Stable latent dynamics in the face of changing neural recordings

Canonical correlation analysis (CCA) effectively identifies linear transformations that maximize the correspondence between two sets of signals. One concern is whether CCA is too powerful, potentially always able to find linear transformations that make empirical latent dynamics look similar, thus misleading us into thinking that the true latent dynamics are stable even if they are not. We argue against this scenario. A fundamental change in the intrinsic dynamics of the full neural population would reduce canonical correlations. Our previous work using CCA provides supporting evidence^12^: when the latent dynamics were shuffled in time and compared using CCA to the original version, the CCs degraded to ≤ 0.3. Additionally, in cases where there is a many-to-few mapping, such as from motor cortex onto the lower-dimensional space of limb muscles, there is not a unique set of dynamics that could generate the observed behavioral output^52^. Simulation work by Sussillo, Shenoy et al. provides such an example. They trained recurrent neural networks (RNNs) to generate outputs that reproduced muscle activity (EMG) recorded during reaching. When they did not constrain the network architecture to be sparsely connected, the CCs between the simulated and actual latent dynamics were low, even though the RNN accurately reproduced the recorded EMGs^30^. This is a compelling example of how a consistent behavior, in this case measured by the EMG output of the RNN, is not necessarily associated with stable latent dynamics. A similar observation of variable latent dynamics leading to consistent behavior has recently been made in the analysis of hippocampal data^52^. On the basis of this evidence, we propose that the stability of latent dynamics reported here is not a trivial consequence of its association with consistent repeated behavior; instead, it reflects a fundamental feature of cortical function that leads to that behavior.

It is a feature of multielectrode array recordings that the exact set of well-discriminated neurons changes progressively over time^17^ (Fig. 3c, Fig. S3). Nonetheless, we have shown that CCA can compensate for the apparent changes in the latent dynamics observed within the changing empirical neural space of the recordings. How is it possible to find stable latent dynamics based on unstable neural recordings? Here we assume that a true low-dimensional neural manifold to which the dynamics of the full neural population is confined does exist^1^ (Fig. 1c). How do we access this true manifold from data? We can apply dimensionality reduction to the empirical neural space spanned by a specific set of recorded neurons. The resulting empirical manifold can only account for the projection of the true manifold onto the neural space of the recordings. As the recorded neurons change, the empirical neural space changes, and so does the projected neural manifold accessible to our measurements (Fig. 1d,e). However, the dynamics within the true neural manifold has unique signatures specific to the task; these get preserved in the projection onto changing empirical neural spaces^11^, making the projected dynamics amenable to alignment. As such, CCA was able to identify linear transformations to align the latent dynamics over days, despite the fact that these latent dynamics had been projected onto different empirical neural spaces. The transformation from the true neural manifold onto the empirical neural manifold involves two linear transformations. First, the true latent dynamics within the full neural space —which incorporates all neurons modulated by the task— is projected onto the empirical neural space of the recorded neurons. This transformation involves a dimensionality reduction from approximately 10^6^ to 10^2^. Then, the empirical latent space embedded within the empirical neural space is found using PCA; this transformation involves a dimensionality reduction from approximately 10^2^ to 10. Since both of these operations are linear, it is thus not surprising that the alignment of empirical latent dynamics can be achieved with a linear method such as CCA.

## Stability of latent dynamics throughout the cerebral cortex

We have reported here on the stable latent dynamics in three cortical areas during execution of the same behavior across days, months, and even years. The activity in each one of these regions is closely tied to the behavior ^50^. We posit that the stability of the behavior and the stability of the corresponding latent dynamics are intimately tied to each other, however, the stability of the population dynamics associated with a specific behavior might well change across brain areas, perhaps decreasing the further removed the brain area is from the periphery and the outside world. For example, decision-making dynamics in prefrontal cortex depend not only on the resulting behavior, but also on internal states (emotion, arousal, etc.) and sensory inputs (sight, sound, touch, etc.) that influence the behavioral decision^53^. There could thus be different neural dynamics that reflect changes in internal state but underlie a stable behavioral output. Within the motor cortex, recent work indicates that the supplementary motor area (SMA) does not exhibit the rotational dynamics consistently observed in M1^54^; note that SMA is a “higher” motor cortical area whose activity reflects motor timing or sequence production^4,55^. We would expect that the dynamics of areas further removed from behavior and more affected by other modulatory or sensory influences are less preserved over long spans of time, as their activity captures more than simply the afferent state of the limb, or observable behavioral output^52^. Still, the results reported here support the hypothesis that when the activity of a brain area is intimately tied to a behavioral output, such as M1 in the context of movement, the underlying latent dynamics corresponding to stable behavior are preserved. Importantly, we have provided evidence that the stability of the latent dynamics is fundamentally of cortical origin, and cannot be trivially explained by non-cortical aspects of the behavior such as the mechanics of movement. For instance, we have shown that latent dynamics in PMd are stable during motor planning, when no behavior has yet occurred.

## Practical implications for braincomputer interfaces

Brain-computer interfaces (BCIs) are based on decoding algorithms that map neural activity onto control variables. This technology holds great promise to revolutionize rehabilitation and assistive technologies^56,57^. Several groups have used BCI decoders based on recorded neural activity to control computer cursors^58,59^, robots^60,61^, and even paralyzed limbs through muscle^62,63^ or spinal cord stimulation^64^. Due to neural turnover, the neural signals that provide inputs to these decoders typically change after a few days, rapidly leading to degraded BCI performance^65,66^. Many groups have been working to develop computational techniques to continually recalibrate these decoders, and thereby restore some of their degraded function^67,68^. Other groups have suggested that the use of input signals such as multiunit threshold crossings^65^ or local field potentials^69^ could reduce the magnitude of these changes, at the risk of reducing the amount of available information. The stable latent dynamics reported here offer an intriguing alternative to existing approaches: BCIs based on latent dynamics as opposed to recorded neural activity could be periodically aligned through a linear procedure such as CCA, thereby achieving stable performance through months or even years of neural turnover. Recent work has shown that approaches based on latent dynamics can be used to improve decoding stability^70,71,38^, adjust for changes in neural inputs^72^, and enable unsupervised decoding^68^.

## Summary

Here we have shown that the latent dynamics associated with the repeated, consistent execution of a given behavior are stable for as long as two years. Our results cover three cortical areas: PMd, M1, and S1. Stability was in each case associated with a different function: movement planning, execution, and somatosensory input processing, respectively. This well-preserved relationship between stable latent dynamics and consistent behavior contrasted with the large changes observed in the activity of single neurons. These observations have broad implications for experiments studying neural activity over time: the activity of individual neurons is best viewed as a sample of underlying true latent dynamics throughout the surrounding brain region. Similar latent dynamics have been identified in many cortical regions (see reviews in Refs.^1,8^), and for a wide variety of tasks, including working memory^73,74^, decision making^13,75,76^, visual^77,78^, olfactory^9,74^, and auditory^79^ discrimination, navigation^80^, and movement^12,14,15,81–83^. Moreover, these latent dynamics exhibit some common characteristics across cortical regions^13,73–75^ and even across species^82,84^. These commonalities suggest that the stabilization of latent dynamics may be ubiquitously exploited by the brain for many behaviorally-relevant purposes, including sensation, perception, decision making, and movement.

The notion of latent dynamics is increasingly accepted as a useful tool for systems neuroscience research, as the low-dimensional neural manifold is more amenable to analysis and visualization than the neural space. It is also a useful tool for brain-computer interfaces design, as a low-dimensional input space facilitates the specification of predictors of movement features. While useful, we posit a view for these latent dynamics that goes beyond that of a mere tool. In this conceptual framework, the computational abilities of the brain are implemented through the dynamics of neural populations; these dynamics are low dimensional and confined to specific neural manifolds; in sensorimotor representations, these manifolds are task-specific and acquired during task learning. Here we have provided evidence that latent dynamics are not just a convenient mathematical abstraction for model building, but the extant and stable building blocks of consistent behavior.

## Acknowledgements

This work was supported in part by “Talent Attraction” Grant 2017-T2/TIC-5263 from the Community of Madrid (J.A.G.), by a grant from the Whitaker International Scholars Program (M.G.P.), and by Grant NS053603 from the National Institute of Neurological Disorder and Stroke (S.A.S. and L.E.M.).

## METHODS

### Behavioral task

We trained six monkeys to sit in a primate chair and make reaching movements using a custom planar manipulandum. All six of the monkeys performed a similar two-dimensional center-out task (Fig. 2a,b) for long periods of time; across all monkeys we collected 137 days of data, with timespans between three weeks and approximately two years. In the task, the monkey moved his hand to the center of the workspace to begin each trial. After a variable waiting period, the monkey was presented with one of eight outer targets (or four targets for Monkey H), equally spaced in a circle and selected randomly with uniform probability. Monkeys C, M, and T were trained to hold during a variable delay period during which the target remained visible before receiving an auditory go cue. Monkeys P and H were not subjected to this delay period. Early recordings from Monkey C also omitted this instructed delay period, though he was later trained on the delayed version of the task. With respect to our main results, we saw no difference between these groups of sessions. To receive a liquid reward, the monkeys were required to reach the outer target within 1 s. Monkeys C, M, and T were required to hold within that outer target for 0.5 s. For Monkey P and early sessions with Monkey H, this outer target hold period was omitted. Monkey H was later subject to a brief hold period of 100 ms, to ensure that he decelerated to end the reach within the target. Thus, there were some kinematic differences between the early and later sessions with Monkey H; since much of the movement was similar, we observed similar results even when all of the recordings and all sessions were considered. As the monkeys performed this task, we recorded the position of the endpoint at a sampling frequency of 1 kHz using encoders in the joints, and digitally logged the specific timing of task events such as the go cue.

### Behavioral data analysis

In all of the following analyses, we considered only the trials in which the monkey successfully achieved the outer target within the specified time and received a reward. We then subselected trials such that all sessions contained an equal number of reaches in each of the target directions. Within each trial, we isolated a window of interest that captured the majority of the movement. Comparison of the dynamics requires each trial on each day to have the same number of data points. We thus adjusted this window slightly according to the behavioral idiosyncrasies (reaching speed, etc.) of each monkey, so as to maximize the number of samples while preserving an equal number of data points across trials. For example, monkeys with naturally slower reach speeds were assigned longer windows. Our results were qualitatively unchanged with reasonable adjustments to this window. For Monkeys M, T, C, and J we used a window starting 120 ms before movement onset and ending 420 ms after movement onset. For Monkeys P and H, we used windows beginning at the go cue and ending after 570 ms or 660 ms, respectively.

We performed all subsequent analyses by comparing all pairs of sessions performed by each monkey, only looking forward in time. For example, with three days of recordings, we compared day 1 to day 2, day 1 to day 3, and day 2 to day 3. In general, when a recording on day *j* was compared to a subsequent recording on day *k*, the result was assigned to (*k* – *j*) days between sessions. First, we studied the hand kinematics to assess behavioral consistency. We took the derivative of the endpoint position to compute the endpoint velocity. Within each session, we ordered all trials by reach direction and concatenated all trials. Since the trials were subselected to equalize the counts across both time and reach directions (last column of Table 1), the resulting data matrices were of equal size for all days. Each matrix entry represented a datapoint from the same time sample and target across all days, allowing for a point-by-point direct comparison of dynamics. To assess the stability of behavior over time, we computed the correlation (Pearson’s *R*) for the 2-D velocity signals between pairs of days in all possible combinations.

### Neural implants

All surgical and experimental procedures were approved by the Institutional Animal Care and Use Committee (IACUC) of Northwestern University. In order to record chronically from populations of cortical neurons, we implanted 96-channel Utah electrode arrays in M1, PMd, or S1 using standard surgical procedures. Monkeys T (male, *macaca fascicularis*) and M (male, *macaca mulatta*) were implanted in PMd of the right hemisphere: Monkey M also received a second array in right M1 in the same procedure. Monkey C (male, *macaca mulatta*) received two implants: first, a single array in right M1 (denoted C_R_ throughout the text), followed years later by implants in both M1 and PMd of the left hemisphere (denoted C_L_). Monkey J (male, *macaca mulatta*) received an array in the left M1. Both Monkeys P and H (male, *macaca mulatta*) received arrays in S1 (Brodmann’s Area 2) of the left hemisphere. Table 1 summarizes the implants and sessions of neural recordings for each monkey.

Neural activity was recorded during the behavior using a Cerebus system (Blackrock Microsystems, Salt Lake City, UT). The recordings on each channel were digitized, band-pass filtered (250-5000 Hz), and then converted to spike times based on threshold crossings. The threshold was set according to the root-mean square (RMS) activity on each channel (Monkeys C, M, T, and J: 5.5×RMS; Monkey P: 4×RMS; Monkey H: 5×RMS). Although most of the analyses of this paper focus on the multiunit threshold crossings on each recording channel, we also manually spike-sorted the recordings from Monkeys M, C, and T to identify putative single-neurons, which we used in the control analyses for M1, as well as in the tracking across days analysis (see below).

### Tracking single neurons over days

For all sessions recorded with Monkeys C, M, and T, we sorted the waveforms that exceeded the threshold to identify putative single neurons. Each of these sorted units can be uniquely described by its waveform shape and its inter-spike interval (ISI) distribution. We applied a statistical test based on these metrics to determine whether or not a given neuron was recorded on two different days^17^. In brief, the waveform shape and ISI of two neurons recorded on a given electrode on two different days was compared against an empirical null distribution taken from neurons recorded on all other electrodes, thus known to be from different neurons. Cells were considered to match if the joint probability of both metrics matching was less than 0.01.

### Analysis of neural spatial tuning across days

We described the spatial tuning of the multiunit on each electrode by using cosine tuning curves^85^. On each recording electrode, we computed the average firing rate within the time window used for decoding or classification (during motor execution for M1 and S1, and during motor planning for PMd, respectively). On each session, we then averaged across all trials for each reach direction and used linear regression to fit the tuning curve according to:

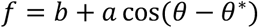

where *b* is the baseline mean firing rate, *a* is the depth of modulation, and *θ\** is the preferred direction^85^; these three parameters describe the directional tuning of the average firing rate *f* for each recording electrode. For each electrode, we tracked the changes in the parameters of this model across all pairs of days for which neural activity was recorded on that electrode on both days. For this subset of electrodes, we assessed the magnitude of the change in mean firing rate, modulation depth, and preferred direction, the latter as a circular difference.

### Neural latent dynamics analysis

To characterize the dynamics of the latent activity associated with the recorded neural activity in each session, we computed a smoothed firing rate as a function of time for the multiunit activity on each electrode. We obtained these smoothed firing rates by applying a Gaussian kernel (s.d.: 50 ms) to the binned square-root-transformed firings (bin size: 30 ms) of each recorded multiunit^10^. We excluded electrodes whose activity had a low mean firing rate (< 1 Hz mean firing rate across all bins), but did not perform any additional preselection, such as based on directional tuning. For each session, this produced a neural data matrix **X** of dimensions *n* by *T*, where *n* is the number of recorded units and *T* is total number of time points from all concatenated trials on a given day; *T* is thus given by: number of targets per day × number of trials per target × number of time points per trial. We performed this concatenation as described above, by subselecting the same number of trials for all sessions and targets for each monkey (Table 1) and ordering the datapoints by time and target. For the analysis of M1 and S1 activity, we considered the same window of interest during a trial as we did for the behavioral analysis (see above); these values were chosen to represent movement execution in M1, and feedback control in S1. For the PMd activity, we analyzed the preparatory activity within a window aligned with movement onset, starting 390 ms before movement onset, and ending 60 ms after movement onset. This window started after the putative visual response elicited by the target presentation^86^, and it was advantageous because it included mostly preparatory activity, but also some of the dynamics of the transition from preparation to movement.

For each session, the activity of *n* recorded multiunits was represented as an empirical neural space, an *n*-dimensional sampling of the state of the cortical area of interest. In this space, the joint recorded activity is represented as a single point whose coordinates are determined by the firing rate of the corresponding multiunits (Fig. 1a). Within this space, we computed the low-dimensional neural manifold by applying principal component analysis (PCA) to the neural data matrix **X**. This yielded *n* principal components (PCs), each a linear combination of the smoothed firing rates of all *n* recorded units. These PCs are ranked based on the amount of neural variance they explain. We defined an *m*-dimensional session-specific neural manifold by keeping only the leading *m* PCs, which we refer to as neural modes (Fig. 1a). Based on previous work by our group and others, we chose the following dimensionality values: *m*=10 for M1, *m*=15 for PMd, *m*=8 for S1. The specific choice of *m* is not critical, as our results held across a broad range of *m* values (Fig. S5a-c).

We computed the latent dynamics within the neural manifold by projecting the time-dependent smoothed firing rates of the recorded neurons onto the PCs that span the neural manifold. This produced a data matrix **L** of dimensions *m* by *T*, where *m* is the dimensionality of the manifold and *T* is total number of time points from all concatenated trials on a given day. Recent theoretical and experimental work has demonstrated that when the recorded neural activity is projected onto the low-dimensional manifold to obtain the latent dynamics, the result is not sensitive to whether the manifold was estimated on the basis of single units or multiunits^33^. Nevertheless, we repeated all the analyses for a subset of three monkeys with M1 implants by using putative single neurons to obtain the manifold, and verified that our results held (Fig. S5d-g).

### Alignment of latent dynamics across days

The substantial turnover across days observed in our neural recordings (Fig. 3, S3,4) implies that the empirical neural space in which the experimentally accessible neural manifold and the latent dynamics are embedded changed across days. Our hypothesis predicts that the true latent dynamics associated with consistent behavior should be stable across days. In order to verify this hypothesis, we need to compensate for the fact that the true latent dynamics is being projected onto different empirical manifolds on different days. If our hypothesis is correct, we expect to be able to compensate for this change in the embedding space by using canonical correlation analysis^34^ (CCA). Given the concatenated single-trial latent dynamics **L**_*A*_ and **L**_*B*_ from two days *A* and *B*, CCA finds linear transformations that applied to **L**_*A*_ and **L**_*B*_ make the corresponding latent dynamics maximally correlated.

CCA starts with a QR decomposition of the transposed latent dynamics matrices **L**_*A*_ and **L**_*B*_, 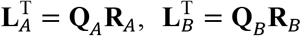. The first *m* column vectors of **Q**_*i*_,*i* = *A,B*, provide an orthonormal basis for the column vectors of 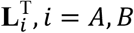. We then construct the *m* by *m* inner product matrix of **Q**_*A*_ and **Q**_*B*_ and perform a singular value decomposition to obtain

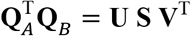

The new manifold directions that CCA finds so as to maximize the pairwise correlations between latent dynamics across the two tasks are the corresponding *m* by *m* matrices

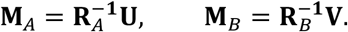

The elements of the diagonal matrix **S** are the resulting canonical correlations (CCs), sorted from largest to smallest. They quantify the similarity in the aligned latent dynamics. For comparison with the CCs between aligned latent dynamics for different days, we computed the CCs between the corresponding unaligned latent dynamics, given by the pairwise correlations between the rows of **L**_*A*_ and **L**_*B*_.

We used the within-day variability in the latent dynamics across blocks of trials for a given day to obtain an upper bound for the across-day CCs. We split all the trials in one day into two nonoverlapping sets of trials, ensuring that the groups were matched by target and time points, and performed CCA on the latent dynamics (100 repetitions). The mean value for each of the *m* ordered CCs in this distribution was used to define the within-day CCs. To represent more compactly how the alignment process compensates for the changes in latent dynamics due to neural turnover and its resultant change in embedding space, we computed the *normalized similarity*, the ratio of the across-day aligned or unaligned CCs to the upper-bound provided by the within-day CCs.

### Decoding hand velocity from motor and somatosensory neural activity

To test whether the aligned latent dynamics in M1 and S1 maintain movement-related information, we built linear decoders to predict the two-dimensional hand velocity from neural data. Our hypothesis was that the aligned latent dynamics should provide accurate predictions of hand kinematics over time. To test this hypothesis, we compared the predictive accuracy of three different types of decoders: 1) a within-day neural decoder trained and tested on the same day based on the recorded neural activity, 2) an across-day neural decoder trained on the neural activity recorded on the first day and tested on neural activity recorded on subsequent days, and 3) an across-day latent decoder trained on the latent dynamics of the first day and tested on the aligned latent dynamics of subsequent days.

All decoders were standard Wiener filters (REFs) that used as inputs the neural activity, either the multiunit firing rates for the within-day and across-day neural decoders or the across-day aligned latent dynamics. We also included three bins of recent spiking history, for a total of 90 ms. These additional neural inputs incorporate information about intrinsic neural dynamics and account for axonal transmission delays. When decoding from M1, whose activity causes the ensuing movement, the additional bins preceded hand velocity signals. When decoding from S1, whose activity is largely in response to the executed movement, the additional bins lagged behind hand velocity signals. The R^2^ value between actual and predicted hand velocity was used to quantify decoder performance as a *predictive accuracy*.

The within-day decoder was trained and tested on the same session, using a six-fold leave-one-out cross-validation procedure to protect against overfitting. Before splitting the recorded neural activity for the session into six blocks, the corresponding trials were shuffled to remove any bias due to time through the session. The R^2^ values for the six test blocks were averaged to obtain a final reported value. The within-day performance provided an upper-bound to the performance of across-day decoders. The across-day neural decoders were computed for all pairwise combinations of days, training on the neural activity recorded on the first of the two days and testing on the later day. The across-day aligned decoders were trained and tested on latent dynamics after alignment. To compare across all sessions and monkeys more easily, we normalized the across-day predictive accuracy by dividing it by the within-day predictive accuracy to obtain the *normalized predictive accuracy*.

### Predicting target direction from PMd planning activity

We trained naive Bayes classifiers to predict the direction of the upcoming movement based on premovement planning activity^20^. As inputs to the classifier we used PMd neural activity recorded during a 450 ms window for all three monkeys (see above). This window focused primarily on the planning period before movement onset. As we did for movement prediction, we trained three types of classifiers: within-day, across-day based on recorded neural activity, and across-day based on aligned latent dynamics. Within the input window, we averaged all activity to obtain a single value representing the activity for that trial, resulting in either an *n*-dimensional neural input or an *m*-dimensional latent input (whether aligned or unaligned).

The naive Bayes classifiers, trained using Matlab (fitcnb), provided a probabilistic assignment of the inputs to one of eight classes, corresponding to the eight possible target directions. The predicted classes were assumed to be independent; hence the classifiers were naive. To ensure that all targets had the same prior probability, the training data included the same number of trials for each target.

Performance of the trained classifiers was quantified by the percentage of correct classifications. To quantify the performance of the within-day classifiers, we performed a cross-validation procedure in which we left out one random trial to each target and trained the classifier on the remaining data. We then tested the classifier on this left-out sample. We repeated this procedure 100 times, and averaged the test performance. As before, we normalized the performance of the across-day classifiers by dividing it by the within-day performance to compute the *normalized accuracy*.

### Statistics

We applied statistical tests to compare the across-day recorded neural activity and across-day aligned latent dynamics. For all statistical tests, we used distributions that had been normalized by dividing by the within-day values, either for canonical correlation or decoding and classification performance. We used a Wilcoxon rank-sum test to compare the distributions with a significance threshold of *P* < 0.001.

**Figure S1.**
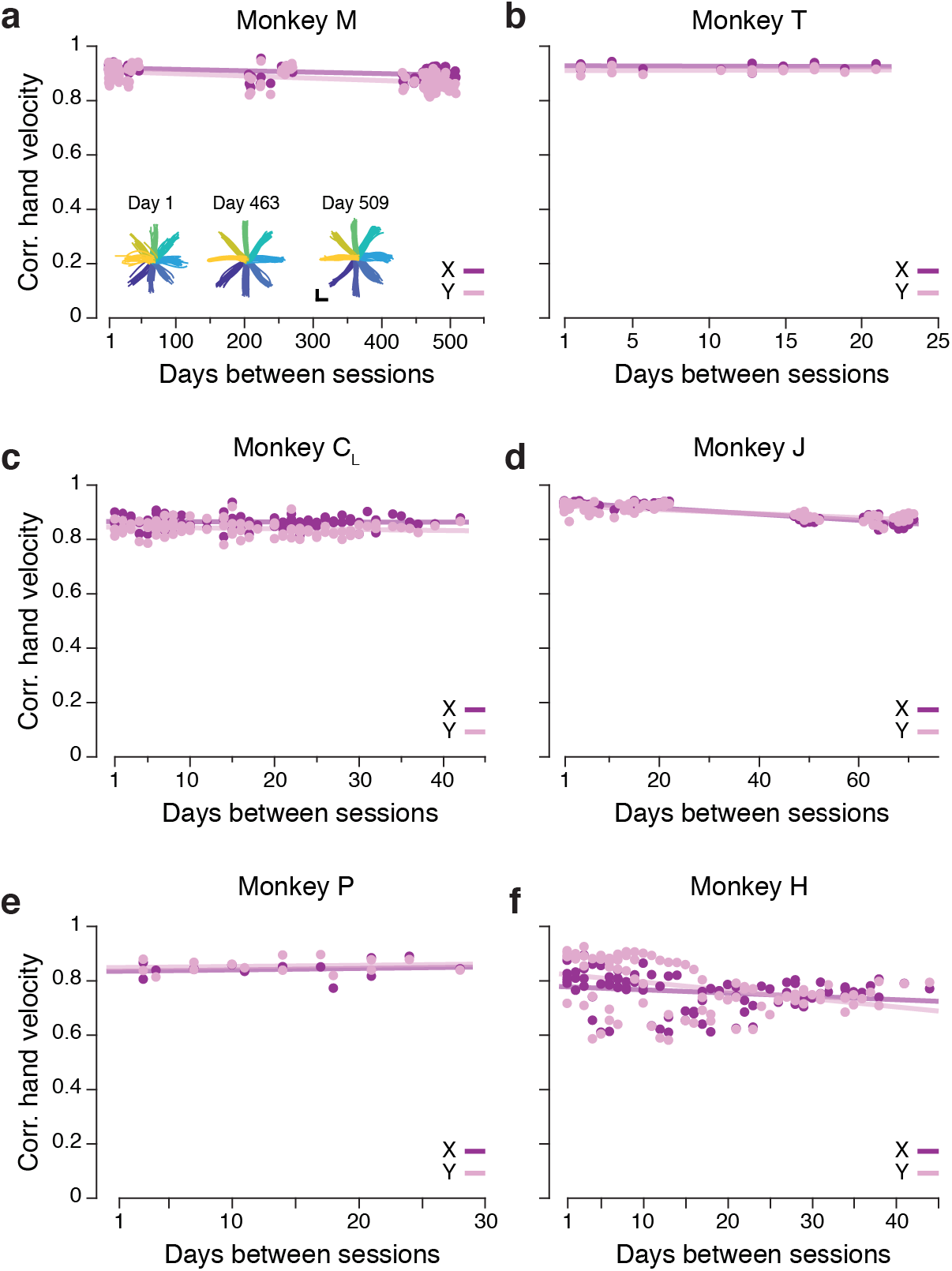
Additional data: task description and consistent behavior. **(a-f)** Correlation between direction-matched single trial X and Y hand velocities across all pairs of days (single dots: individual trials; lines: linear fits) from Monkey M (a), Monkey T (b), Monkey C_L_ (c), Monkey J (d), Monkey P (e), and Monkey H (f).

**Figure S2.**
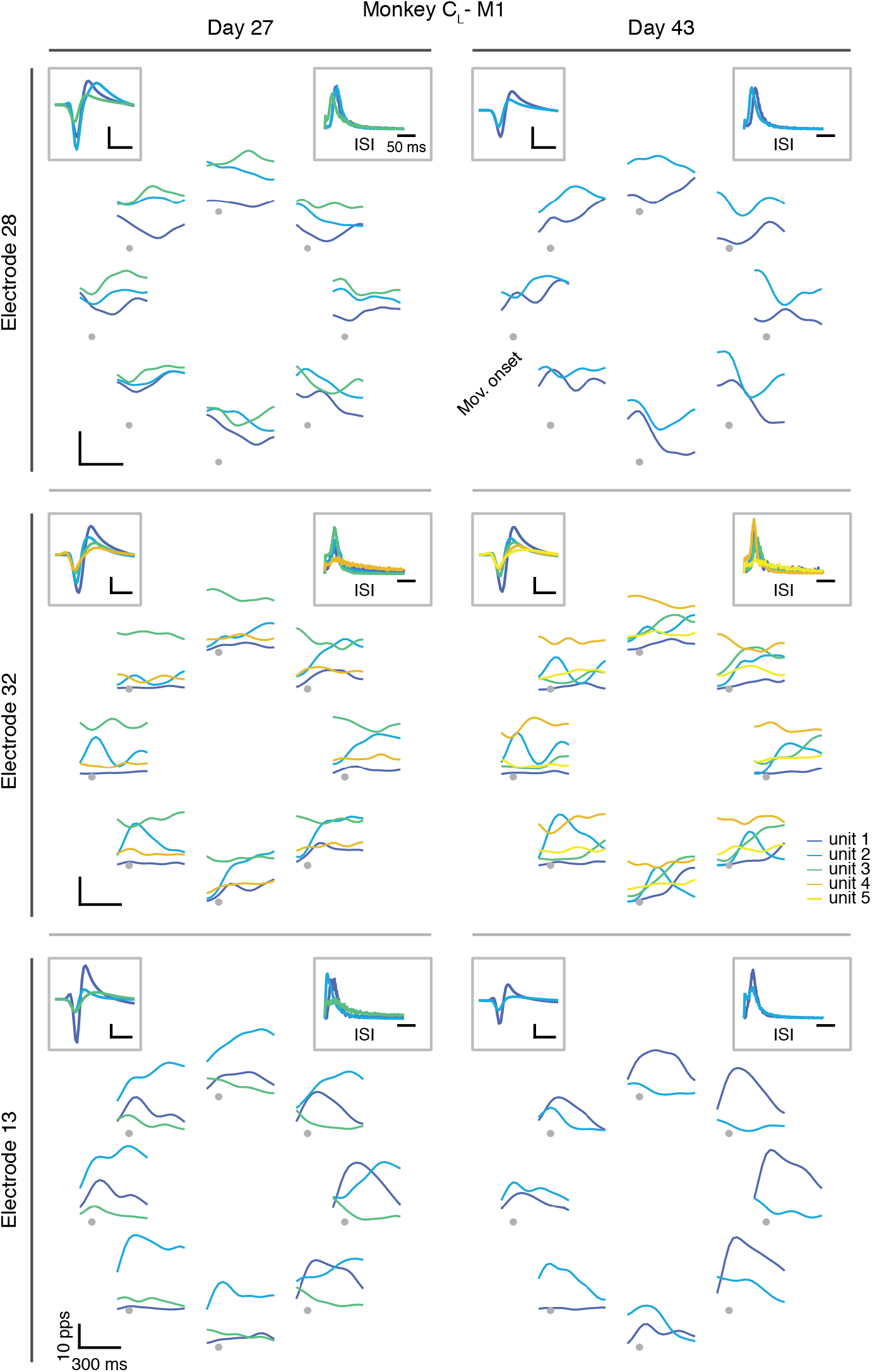
Example neural activity during reaching on two days from Monkey C_R_. Each row shows the firing rates on a different electrode for Day 27 (left column) and Day 43 (right column). Each color represents a different sorted neuron. The eight plots arranged in a circular manner show the firing rate as a function of time during a reach to each of the eight targets, aligned on movement onset and averaged across all trials to the same target. The inset in the top left of each panel shows the average waveform of each sorted neuron; the inset at the top right shows the ISI distribution for each sorted neuron.

**Figure S3.**
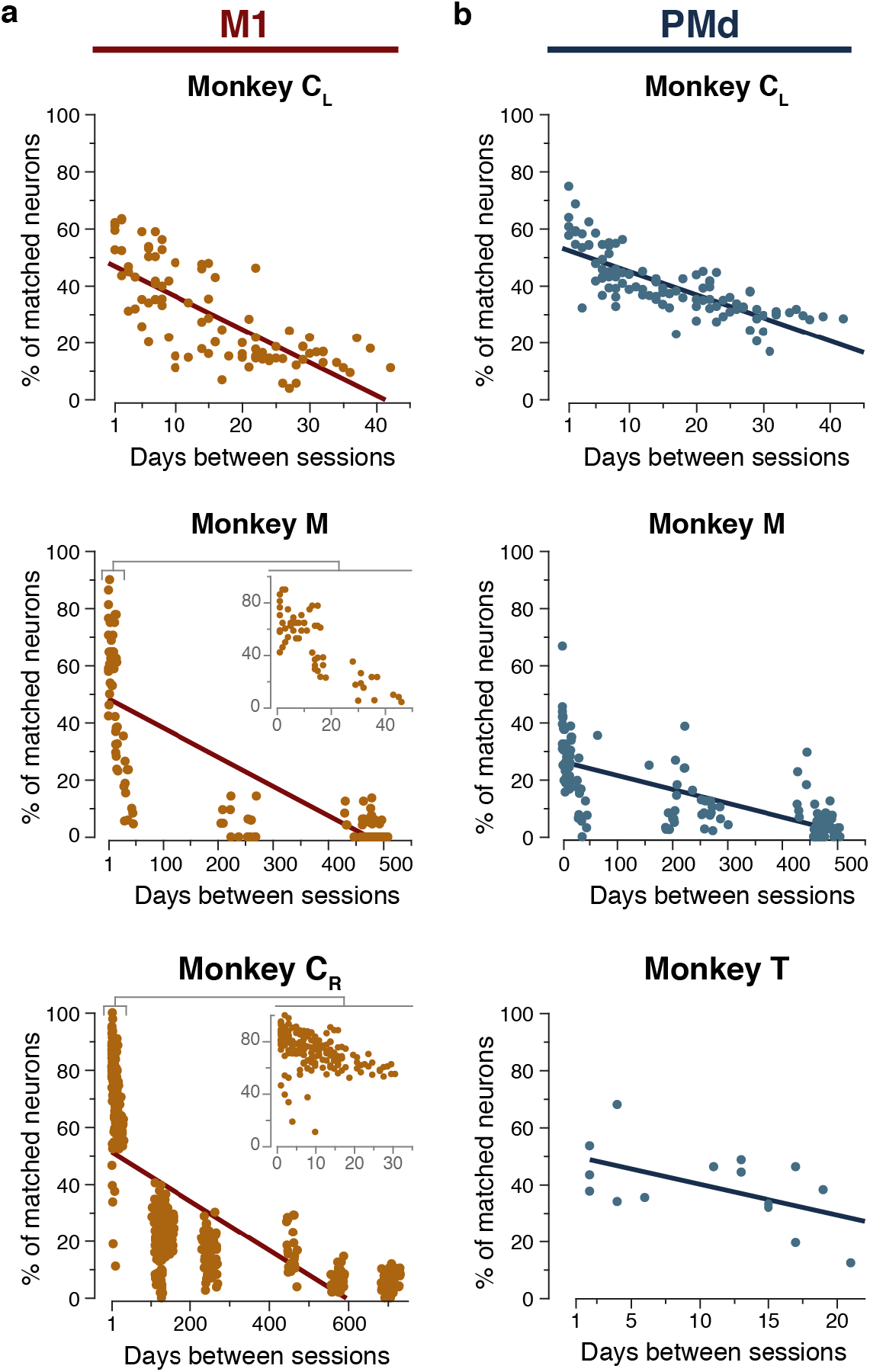
Additional data: neural recording stability. **(a)** Percentage of individual sorted M1 neurons matched across all pairs of days based on action potential waveforms and inter-spike interval (ISI) histograms. Data from Monkey C_L_ (top; duplicated from Fig. 3d), Monkey M (middle; inset highlights the first 50 days), and Monkey C_R_ (bottom; inset highlights the first 35 days). **(b)** Percentage of individual sorted PMd neurons matched across all pairs of days as in (a). Data from Monkey C_L_ (top), Monkey M (middle), and Monkey T (bottom).

**Figure S4.**
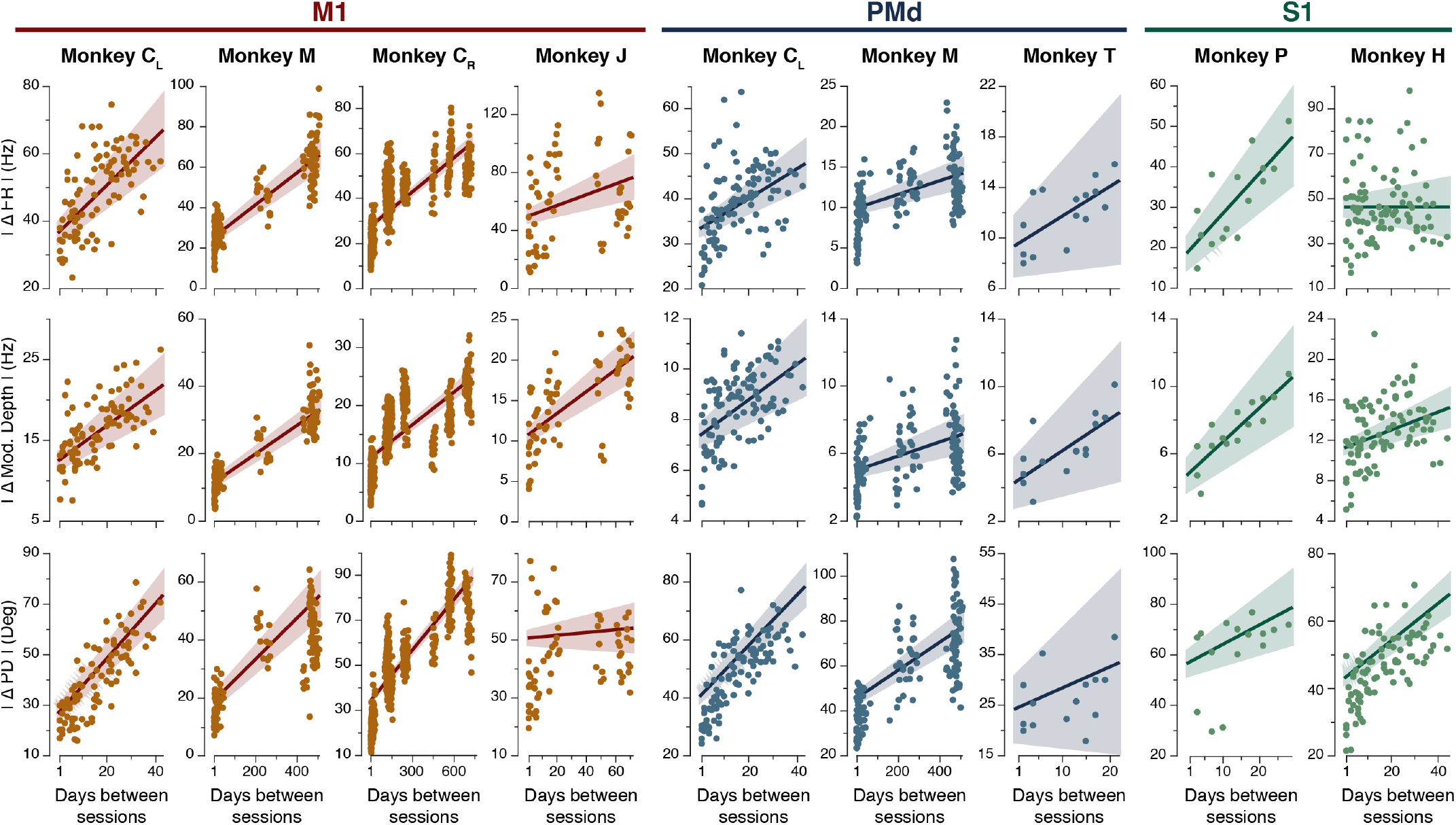
Additional data: neural tuning stability. For each implant: change in mean firing rate (top plot), modulation depth (middle plot), and preferred direction (bottom plot) of standard cosine tuning fits to multiunit activity across all pairs of days. Line and shaded areas: mean ± s.e.m. Plots are grouped by implant and brain area (M1: left; PMd: middle; S1: right).

**Figure S5.**
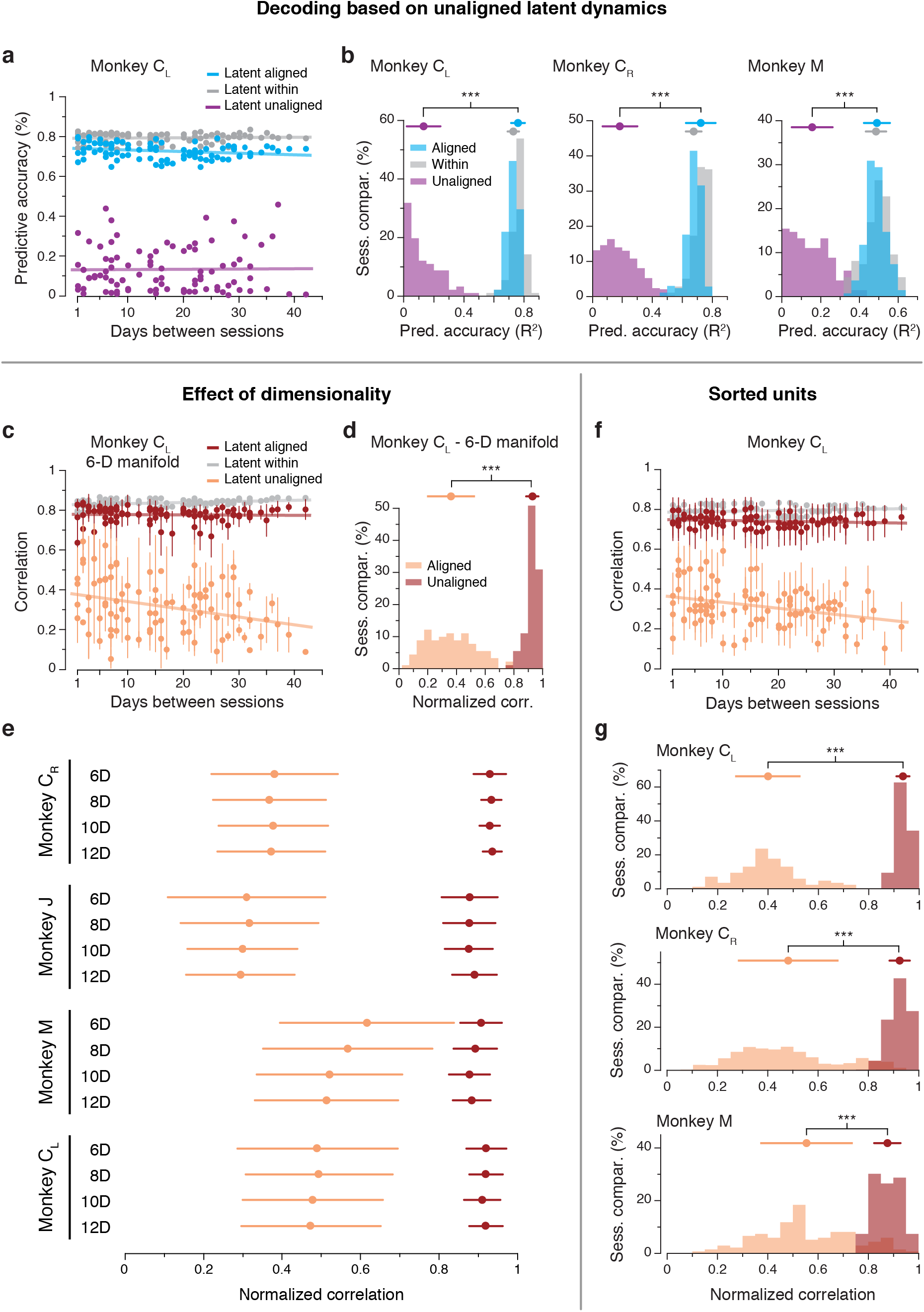
Controls for the alignment procedure using M1 data. **(a)** Predictive accuracy when decoding hand velocity for all pairs of days from Monkey C_L_, using the unaligned latent dynamics as inputs instead of the multiunit activity used in Figure 4. **(b)** Predictive accuracy of decoders using the latent dynamics within-day and across-day (unaligned), as well as across-day after alignment, for Monkeys C_L_, C_R_, and M. **(c)** Correlation of the M1 latent dynamics averaged over the top four neural modes across all pairs of days from Monkey C_L_ using a 6-D manifold (single dots: pairs of days; lines: linear fits). **(d)** Normalized similarity of the aligned and unaligned M1 latent dynamics in the 6-D empirical latent space for Monkey C_L_. **(e)** Mean and s.e.m. for normalized similarity distributions as shown in (b), for all four M1 implants for 6, 8, 10, and 12-D manifolds. The 10-D data presented here summarizes the distributions shown in Figure 4. The significance of the separation between aligned and unaligned distributions held regardless of the choice of latent space dimensionality. **(f)** Correlation of the M1 latent dynamics averaged over the top four neural modes across all pairs of days from Monkey C_L_ using sorted neurons rather than multiunit activity (single dots: pairs of days; lines: linear fits). **(g)** Normalized similarity of the aligned and unaligned M1 latent dynamics in the standard 10-D empirical space obtained using sorted neurons for Monkeys C_L_, C_R_, and M. Error bars: Mean ± s.d.

**Figure S6.**
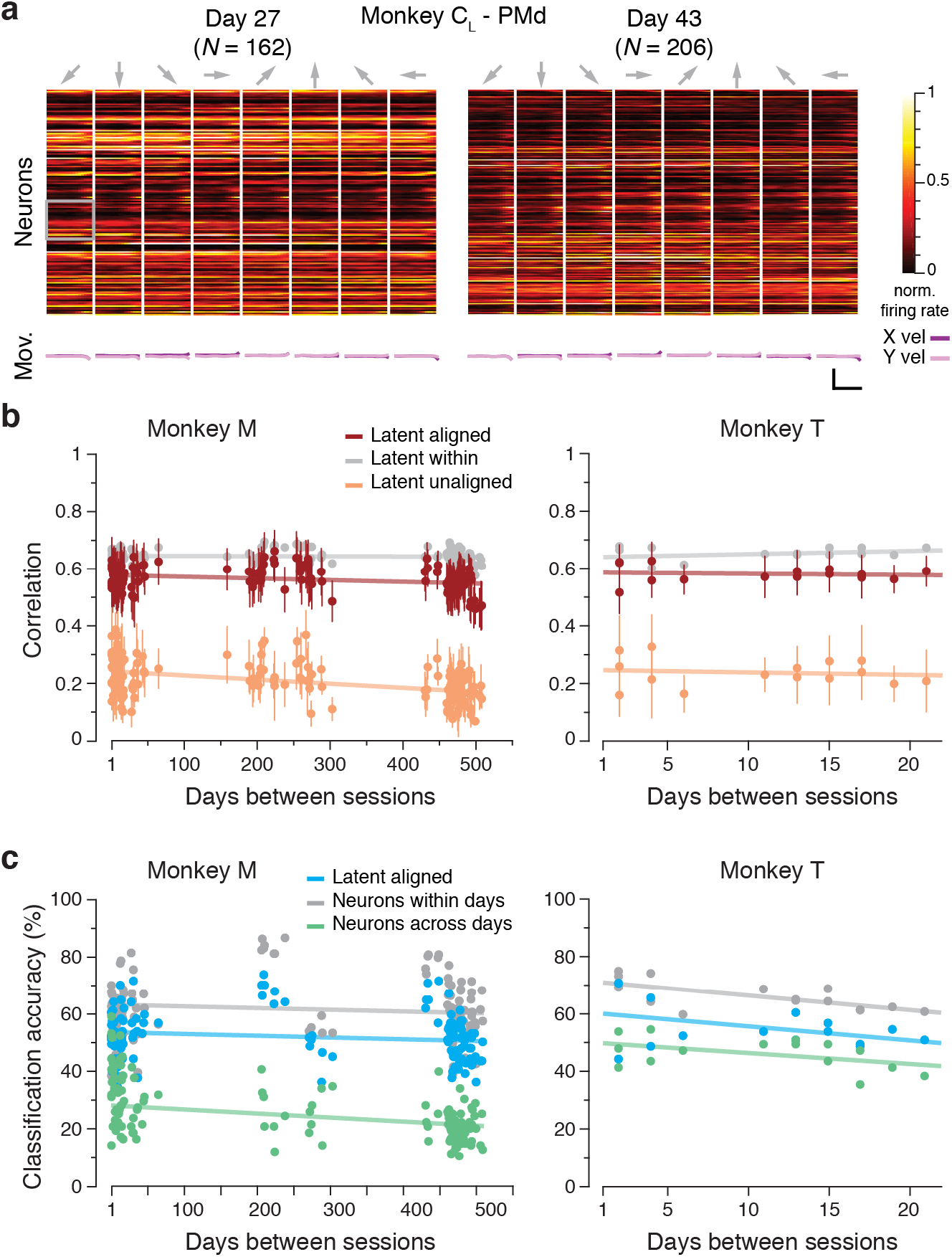
Additional data: PMd alignment and decoding. **(a)** Peristimulus time histograms for all sorted PMd neurons identified on Day 27 and Day 43 from Monkey C_L_ (top; each neuron shown in a different row) and corresponding hand velocity (bottom). Each column represents the average of all trials to each of the eight reach directions (indicated by the arrows above each column). Data was recorded during the pre-movement planning and the transition to movement; hand velocities are thus largely zero. Note the substantial changes in the planning activity of the recorded PMd neurons across days. **(b)** Correlation of the PMd latent dynamics averaged over the top four neural modes across all pairs of days from Monkey M (left) and Monkey T (right) (single dots: pairs of days; lines: linear fits). Error bars: Mean ± s.d. **(c)** Classification accuracy for models trained and tested on all different pairs of days for Monkey M (left) and Monkey T (right).

**Figure S7.**
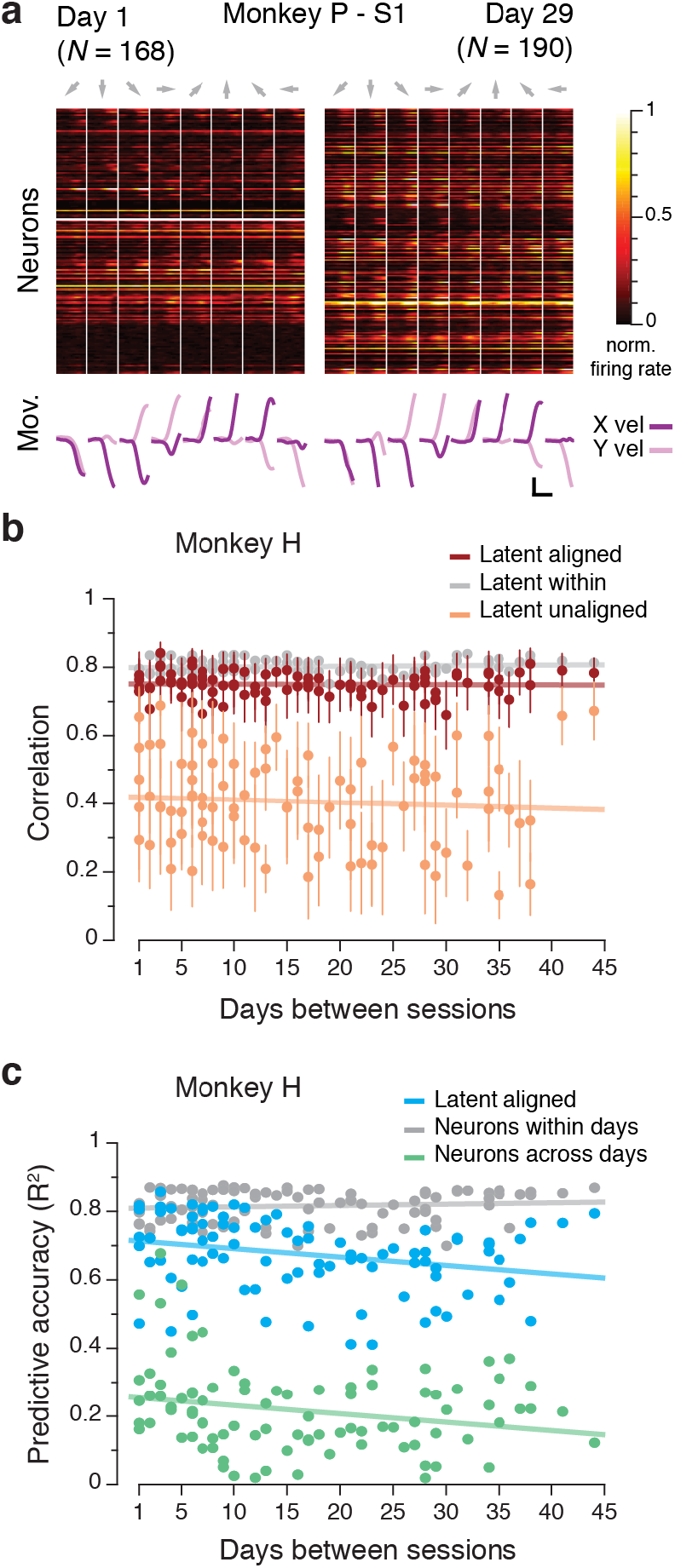
Additional data: S1 alignment and decoding. **(a)** Peristimulus time histograms for all sorted S1 neurons identified on Day 1 and Day 29 from Monkey P (top; each neuron shown in a different row) and corresponding hand velocity (bottom). Each column represents the average of all trials to each of the eight reach directions (indicated by the arrows above each column). **(b)** Correlation of the S1 latent dynamics averaged over the top four neural modes across all pairs of days from Monkey H (single dots: pairs of days; lines: linear fits). **(c)** Predictive accuracy for models trained and tested on all different pairs of days for Monkey H.

